# Pervanadate-induced oxidation relieves autoinhibition of SRC protein tyrosine kinase

**DOI:** 10.1101/2025.04.27.650842

**Authors:** Katie E. Mulholland, Maxime Bourguet, Nuo Cheng, Oisharja Rahman, Daria Ezeriņa, Leonard A. Daly, Tiffany Lai, Silvia Aldaz Casanova, Pau Creixell, Claire E. Eyers, Joris Messens, Patrick A. Eyers, Dominic P. Byrne, Hayley J. Sharpe

## Abstract

Dynamic regulation of protein tyrosine phosphorylation (pTyr) by phosphatases (PTPs) and kinases enables cells to sense and respond to environmental changes. The widely used chemical probe Pervanadate (PV) induces accumulation of high levels of pTyr in cells, an effect primarily attributed to its properties as a PTP inhibitor. This led to the assertion that PTPs are the master gatekeeper of intracellular pTyr homeostasis. Here, we use diverse approaches to reveal that PV disrupts cellular redox homeostasis and directly activates SRC family tyrosine kinases *via* oxidation of specific cysteine residues. Using mass spectrometry and biophysical approaches, we show that oxidation activates SRC by disrupting autoinhibition and altering phosphopeptide binding by its SH2 domain. We further establish that redox-sensitive cysteine residues are essential for SRC to promote cellular overgrowth. Our findings call for a re-evaluation of PV-based experiments and provide compelling evidence that oxidation is a crucial mechanism in controlling the oncogenic properties of SRC.

## Introduction

Phosphorylation of protein tyrosine residues (pTyr) is a critical posttranslational modification that orchestrates diverse cellular processes (*1*) and is often dysregulated in disease (*2, 3*). Dynamic pTyr levels are controlled by the activities of protein tyrosine phosphatases (PTPs) and kinases (PTKs). Disruption of both PTPs and PTKs can lead to pathological conditions, highlighting the importance of understanding their regulation in homeostasis and disease. The development of specific PTP inhibitors has been challenging, with only a few compounds advancing to clinical trials (*4, 5*). As a result, generic PTP inhibitors remain invaluable in gaining insight into pTyr functions and regulation. Of these, orthovanadate (OV), commonly added to cell lysis buffers, and pervanadate (PV), are important tools (*6*). OV acts as a phosphate mimetic that competitively inhibits PTPs, while PV is produced by combining hydrogen peroxide (H_2_O_2_) with OV, followed by the addition of catalase to remove excess H_2_O_2_ (**Fig. 1A**) (*6, 7*).

**Fig. 1.**
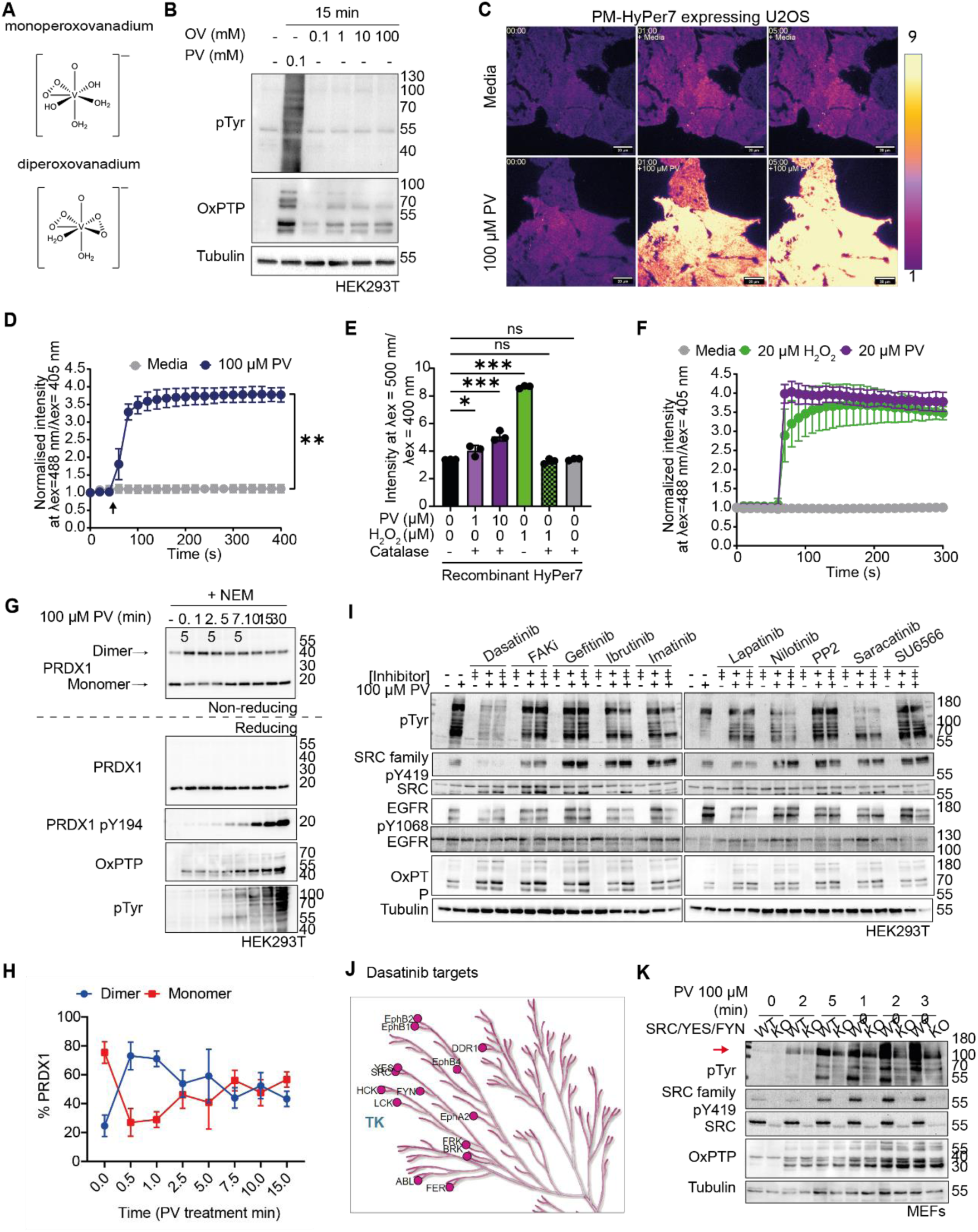
PV perturbs redox homeostasis and induces tyrosine phosphorylation in SRC dependent manner. **(A)** Schematic of mono-peroxo and di-peroxo-vanadium (PV compounds) generated using ChemDraw. **(B)** HEK293T cells were treated as shown, lysed and subjected to immunoblot analysis using the indicated antibodies. Image representative of N=3 experiments. (**C**) Representative false colour images of F488/405 ratio (scaled 1-9) from **movies S1 and S2** of U2OS cells stably expressing plasma membrane (PM) localised HyPer7 and treated with 100 μM PV or media. Scale bar = 20 μm. (**D**) Quantification of the ratio of the two excitation maxima (F488/405), normalized against initial value, of PM-HyPer7 expressing U2OS cells over a time course, imaged every 20 s. Treatments were added at 60 s (arrow). Mean of n=3 ± SD plotted. ** p ≤ 0.001 Unpaired t-test at t= 400 s. (**E**) Ratio of the two excitation maxima (F500/400) of recombinantly expressed HyPer7 (2.5 μΜ) following its treatment with the indicated reagents. The bars represent the mean of n=3 ± SD. *** p ≤ 0.001; ns – non-significant. The p-value is based on Dunnett’s multiple comparisons test of a one-way ANOVA. (**F**) Quantification of the ratio of the two excitation maxima (F488/405), normalized against initial value, of PM-HyPer7 expressing U2OS cells over a time course, imaged every 20 s. Treatments were added at 60 s (arrow). Mean of n=3 ± SD plotted. (**G**) Cells were subjected to the indicated PV time course, washed with PBS with catalase and NEM, prior to lysis without NEM. Samples were resolved on non-reducing or reducing gels as indicated to distinguish PRDX1 monomers from disulfide-linked dimers. Total and phospho PRDX1 blots are from separate gels but the same samples. Representative of n=3 experiments. (**H**) Densitometric quantification of PRDX1 dimer and monomer bands from G. Mean of n=3 ± SD plotted. (**I**) HEK293T cells were treated with the indicated tyrosine kinase inhibitors at IC_50_ (+) and 2.5x IC_50_ (‡) concentrations for 2 h prior to addition of 100 µM PV for 15 min before lysis and immunoblotting with annotated antibodies. (**J**) Dasatinib targets based on (*71*) and mapped onto the tyrosine kinase arm of the Human kinome using kinmap (http://kinhub.org/kinmap/). (**K**) WT and Src/Yes/Fyn (SYF) WT and KO cells were stimulated with 100 µM PV for the indicated times prior to lysis and immunoblotting with indicated antibodies. Red arrow indicates 130 kDa. Representative of 3 experiments. See also Figs. S1-S3.

PV exists in multiple oxidation states, containing up to three peroxogroups (*8, 9*). In cells, PV treatment rapidly elevates pTyr levels, enabling PTP substrates to be analyzed through ‘trapping’ and dephosphorylation approaches (*10*). PV is established to inhibit PTPs by hyperoxidizing the low pKa catalytic cysteine to sulfonic acid (*11*). Due to the structural similarity between phosphate and vanadate, it has been proposed that PV exhibits high affinity for the PTP active site (*6*), leading to the conclusion that its effects are predominantly driven through PTP inhibition (*12*). Moreover, the capacity of PV to reproduce key cellular activation states has fostered the prevailing view that PTPs are constitutively active, constantly removing pTyr from proteins (*13*). Furthermore, this concept is a key pillar in the kinetic segregation model, which proposes that steric exclusion of the highly abundant receptor PTP, CD45, is sufficient to trigger T cell activation (*14*), based on experiments showing that PV mimics T cell activation (*15, 16*).

Cellular effects of PV are often indirectly linked to PTK activation, since PTPs are assumed to suppress activity (*12*). However, PTP function is more complex, with several PTPs, including CD45, preventing autoinhibition of PTKs via inhibitory phosphosites (*17, 18*). Furthermore, the catalytic activities of certain PTKs, including EGFR and SRC, are enhanced following reversible oxidation of conserved regulatory cysteine residues (*19-24*). Given that PV possesses peroxo groups with oxidizing potential, we hypothesized that its cellular effects on pTyr levels might extend beyond PTP inhibition to include direct oxidation of redox-sensitive cysteines in other proteins (*25, 26*), including PTKs. Here, we show that the SFKs SRC and YES1 are directly activated by PV. Using conformational analysis and biophysical approaches, we have determined the mechanisms by which PV-induced oxidation relieves SRC autoinhibition. Additionally, we identified a key role for redox-active cysteines in SRC-regulated cell growth and control of Hippo signaling. Our study defines redox-based mechanisms for SRC activation and highlights the importance of cysteine oxidation in oncogenic transformation. In addition, our data call for a mechanistic reinterpretation of the effects of pervanadate on phosphotyrosine and redox proteomes.

## Results

### Pervanadate promotes cysteine thiol oxidation

We initially set out to characterize cellular effects of the PTP inhibitors PV and OV. As expected, PV induced a rapid pTyr accumulation in HEK293T cells with concomitant irreversible PTP oxidation (OxPTP) (**Fig. 1B**). In contrast, the PTP inhibitor OV had minimal effect on global pTyr despite the induction of irreversible PTP oxidation at higher doses (**Fig. 1B**). Longer treatment (4h) with high doses of OV, resulted in more pronounced upregulation of pTyr and PTP hyperoxidation, consistent with previous reports of oxidative damage by OV (*27*) (**Fig. S1A**). We next asked whether our OV preparations inhibit PTPs. First, we confirmed that OV inhibits the phosphatase activity of recombinant purified PTPRK *in vitro*, albeit less potently than PV (**Figs. S1B-S1C**), aligning with previous reports (*28*). Furthermore, Flag-PTPRK immunoprecipitated from OV or PV treated cells was catalytically inactive (**Figs. S1D-S1E**). Importantly, and in contrast to PV, OV did not induce extensive tyrosine phosphorylation at this concentration (**Figs. S1F-S1G**). Collectively these data suggest that PTP inhibition alone is necessary, but not sufficient, for high-level global induction of pTyr. We next explored the possibility that PV could have functions beyond PTP inhibition that promote supraphysiological levels of pTyr accumulation.

Since the pTyr-inducing function of PV is usually attributed to PTP oxidation, we investigated whether it has broader oxidizing activities in cells. To this end, we used HyPer7, a ratiometric biosensor that is oxidized by peroxides and reduced by thioredoxins and reduced glutathione (GSH) within cells (*29-31*). Treating U2OS cells expressing myristoylated, plasma membrane-anchored HyPer7 with 100 µM PV led to rapid biosensor oxidation compared to media-treated cells (**Figs. 1C-1D and movies S1-S2**). In contrast, treatment of cells with 50-500 fold higher levels of OV decreased the ratio of oxidized:reduced HyPer7 compared to media alone (**Fig. S1H**). *In vitro*, PV directly oxidized purified recombinant HyPer7, although far less potently than H_2_O_2_ (**Fig. 1E**). By comparison, both 20 µM PV and H_2_O_2_ rapidly oxidized HyPer7 in cells equivalently (**Fig. 1F**). Thus, despite lower direct reactivity with HyPer7, PV is as effective as H_2_O_2_ (at 20 µM) at inducing HyPer7 oxidation in cells. This indicates that PV elicits its effects on HyPer7 in cells by perturbing antioxidant pathways, such as the thioredoxin pathway.

We next performed a time course of PV treatment and monitored the oxidation state of peroxiredoxin 1 (PRDX1), an endogenous substrate of the thioredoxin pathway. Upon oxidation by H_2_O_2_, PRDX forms disulfide-linked dimers, which are detectable as slower migrating species after non-reducing SDS-PAGE. To block post-lysis oxidation, samples were treated with catalase and N-ethyl maleimide (NEM). Within 30s, PV increased PRDX1 disulfide-linked dimers that were undetectable under reducing conditions (**Figs. 1G, 1H and S1I**). We also detected a time dependent increase in PRDX1 phosphorylation (pY194), which is associated with impaired PRDX catalytic function (*32*). Together these data suggest that PV fundamentally perturbs redox homeostasis. Strikingly, PTPs were irreversibly oxidized 30 s post PV treatment, several minutes before the peak in global tyrosine phosphorylation is evident, suggesting sustained or induced PTK activity. Since several kinases also possess redox-active cysteines (*19-24*), we next focused on the role of PTKs in driving the PV pTyr response.

### PV-induced pTyr depends on specific PTKs, including SRC

To determine which PTKs might contribute to PV-induced increases in pTyr, we took an unbiased approach pre-treating cells with a panel of pharmacological PTK inhibitors. PV was equally effective at oxidizing PTPs in cells treated with the inhibitors but dasatinib and saracatanib profoundly limited PV-induced pTyr levels (**Figs. 1I and S2**). Dasatinib and saracatanib inhibit SFKs, ABL and EPH kinases (**Fig. 1J**). SRC activation has previously been linked to PV-induced pTyr signaling (*12, 33*), so we employed genetic manipulation to validate the role of SFKs in this process. We observed an attenuated pTyr response when treating Src/Yes/Fyn knockout (KO) MEFs compared to WT cells when treated with 100 µM PV, despite equivalent levels of PTP oxidation (**Fig. 1K**). We also detected increased Abl1 expression in KO MEFs (**Fig. S3A**), which could explain the hyperphosphorylated protein band at ∼130 kDa induced by PV (**Fig. 1K, red arrow**). However, knockdown of Abl1 in SYF KO MEFs did not further attenuate the PV response or affect the intensity of the 130 kDa band (**Fig. S3B**). Knocking out SRC in HAP1 cells also decreased the levels of pTyr induced in response to PV (**Fig. S3C**). Additionally, knockdown of SRC and YES1, but not FGR, in HEK293T cells led to reduced PV stimulated pTyr (**Fig. S3D**). Thus, SFKs are key mediators of the cellular PV response.

### Pervanadate activates SRC via cysteine oxidation *in vitro* and in cells

We hypothesized that PV induces potent pTyr accumulation in cells by directly activating SFKs. We focused our initial experiments on SRC since its redox sensing via specific cysteines is the best characterised amongst SFKs (*20*). Using purified recombinant SRC, we determined that PV dose-dependently increased SRC kinase activity *in vitro* (**Fig. 2A**). Introducing a pre-incubation period increased activation at lower PV concentrations but decreased catalytic activity at higher PV concentrations (**Figs. 2A and S3E**). OV treatment had no effect on SRC activation (**Fig. 2A**). Similar activity profiles have been reported for SRC and EGFR in cells treated with H_2_O_2_ (*21, 22*), where higher peroxide concentrations were inhibitory potentially due to oxidation of additional cysteines. In support of a redox mechanism, we also observed H_2_O_2_ stimulation of SRC activity, but only at higher concentrations than PV (**Figs. 2B and S3E**). To test whether PV induces cellular SRC oxidation, we used differential alkylation to reveal TCEP-reversible thiol modifications by biotin labelling. At 0-250 µM PV, SRC and the PTP SHP2 were increasingly enriched by streptavidin, indicating increased reversible oxidation (**Fig. 2C**). Higher PV concentrations decreased streptavidin-enrichment of both proteins, suggesting thiols had either become reduced, or irreversibly oxidized and therefore resistant to reduction and alkylation. In fact, numerous biotinylated, reversibly oxidized, proteins showed this trend. Additionally, we observed oxidation of both SRC and SHP2 in the absence of PV, indicating a basal level of oxidative signaling in HEK293T cells, consistent with the presence of PRDX1 dimers in untreated cells (**Fig. 1G**). Together, these data demonstrate reversible oxidation of SRC by PV in cells.

**Fig. 2.**
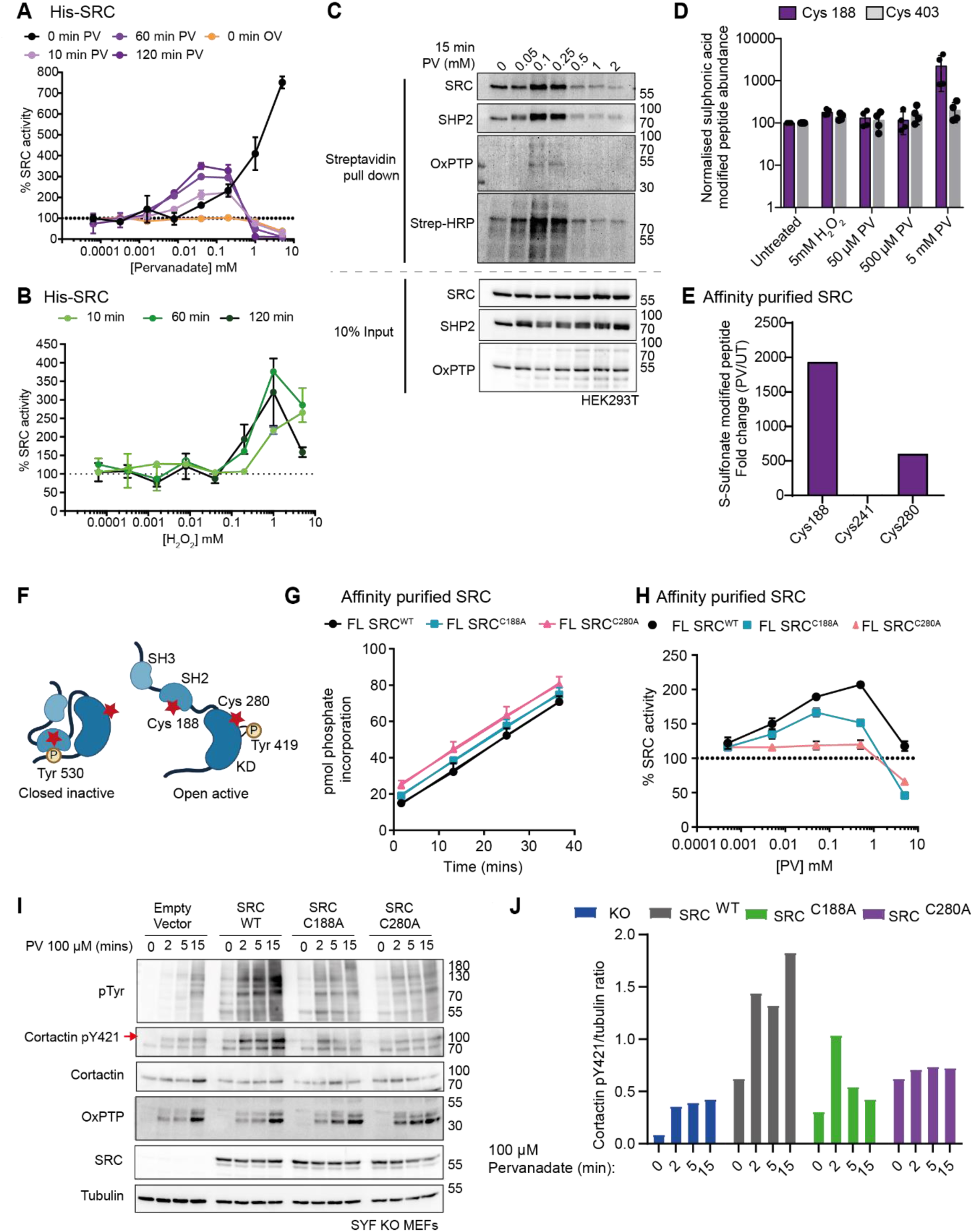
SRC is oxidized on specific cysteines and activated by PV. (**A**) 6xHis-SRC (3 ng) dependent phosphorylation of a specific fluorescent substrate (2 µM). Data shown is relative SRC activity in the presence of the indicated concentrations of PV compared to a buffer only control. SRC was preincubated for 0, 10, 60 or 120 mins prior to initiation of the kinase assay. Means ± SD n=3 experiments are shown. (**B**) Dose response curve for H_2_O_2_ pre-incubated with recombinant 6xHis-SRC (3 ng) for the indicated time periods. Reactions were initiated with the addition of peptide substrate and ATP, and phosphorylation was measured after 15- and 25-mins assay time. Data shown is relative SRC activity in the presence of the indicated concentrations of H_2_O_2_ compared to a buffer only control. Means of n=3 ± SD shown. (**C**) HEK293T cells treated with the indicated concentrations of PV for 15 min, were lysed in catalase and NEM-containing buffer. Reversibly oxidised thiols were reduced with TCEP, and newly regenerated thiol(ate)s were labelled with Iodoacetyl-PEG2-Biotin. Lysates and biotinylated proteins were then affinity purified using streptavidin beads and subjected to immunoblotting with indicated antibodies. Blot representative of 3 experiments. (**D**) HEK293T cells transiently expressing StrepII-tag fused SRC were treated as indicated, and SRC affinity purified by streptactin pull down. Purified protein was then subjected to tryptic digest and analysed by LC/MS-MS for triply oxidised cysteine (sulphonic acid). (**E**) SRC isolated from transfected HEK293T cells by streptactin affinity purification was incubated with or without 5 mM PV, followed by tryptic digest and analysis by LC MS/MS. Values are normalized against abundance of non-cysteine containing SRC peptides. (**F**) Schematic of SRC kinase indicating the position of two key regulatory phosphosites (Circle with P) as well as the two most readily oxidixed cysteine residues (red stars). (**G**) Surface electrostatic potential (*72*) for regions surrounding Cys 280 (PDB:2SRC) and Cys 188 and Cys241 (PDB:1Y57). Units = K_b_T/e_c_. (**H**) Phosphorylation of the fluorescent peptide substrate by WT (black), C188A (blue) and C280A (red) SRC isolated from HEK293T cells by streptactin affinity purification was measured after 30 min. Means of n=3 ± SD are shown. (**I**) Dose-response curves for PV. WT (black), C188A (blue) and C280A (red) SRC isolated from HEK-293T cells by streptactin affinity purification and incubated with the indicated concentrations of PV for 10 mins. Phosphorylation of the fluorescent peptide substrate was measured after 15 min. Data shown is % SRC activity compared to a buffer control. Means of n=3 ±SD are shown. FL = full-length. (**J**) SYF KO MEFs were retrovirally-transduced with specified SRC expression constructs and treated with PV as shown, before lysis and immunoblotting with indicated antibodies. Red arrow indicates relevant band. Representative of >3 experiments. (**K**) Densitometric quantification of western blot in Fig. 2J. Ratios of phosphorylated cortactin/total tubulin levels are shown. See also Fig. S3.

To gain insight into the molecular mechanism(s) of SRC oxidation, we used a targeted approach to identify cysteines oxidized by PV. At high PV concentrations, SRC cysteine thiols appeared to be hyperoxidized to sulfinic or sulfonic acid (**Fig. 2C**), which are identifiable by mass spectrometry (MS) without the need for differential alkylation. We therefore stimulated cells overexpressing SRC with 5 mM PV for 30 min and analyzed affinity purified protein by MS (**Fig. 2D and Table S1**). Peptides containing Cys 188 and Cys 403 were identified, however, only Cys 188 showed a large increase in sulfonic acid modification following PV treatment. We next analyzed purified, untagged SRC treated with PV, which identified sulfonic acid modification of cysteines 188, 280 and 241 (**Fig. 2E and Table S2**). SRC Cys 241 was less sensitive to hyperoxidation by PV than Cys 188 and Cys 280 (**Fig. 2E**), indicating distinct accessibility or reactivity of individual cysteines (*34*). SRC comprises N-terminal, SRC homology (SH)3, SH2 and kinase domains (**Fig. 2F**). Cys 188 is located within the SH2 domain pTyr binding pocket and is unique to SRC amongst SFKs, whilst Cys 280 resides in the kinase domain P-loop of SRC, YES1 and FGR (**Figs. 2F and S3F**). Both Cys 280 and Cys 188 are found close to or within positively charged pockets on the surface of SRC (**Fig. 2G**). In contrast, Cys 241, which was not oxidized by PV, is surface exposed in an electrostatically neutral region of the SH2 domain (**Fig. 2G**). Together this reveals selective oxidation of Cys 188 and 280 by PV.

The conservation of Cys 280 in SFKs prompted us to also evaluate sensitivity of YES1 activity to oxidation, especially given its role in the PV response in cells (**Fig. S3D**). Like SRC, PV dose dependently increased YES1 activity (**Fig. S3G**). In contrast, H_2_O_2_ had a negligible impact on activity (**Fig. S3H**), indicating differences in sensitivity to each oxidant. To directly assess the impact of Cys 188 and Cys 280 on SRC kinase activity *in vitro*, we affinity purified untagged WT (SRC^WT^) and mutant (SRC^C188A^ and SRC^C280A^) forms. All three proteins displayed equivalent kinase activity (**Fig. 2H**). However, when treated with PV, a dose-dependent increase in SRC^WT^ activity was evident, which was absent for SRC^C280A^ and impaired for SRC^C188A^ (**Fig. 2I**). At higher PV concentrations, all proteins were inhibited, potentially indicating oxidation of additional cysteines. Thus, PV-induced SRC activation requires Cys 188 and Cys 280.

We next asked whether these cysteine residues are important for the cellular pTyr response to PV. To this end, we introduced chicken Src and mutants equivalent to C188A and C280A into SYF KO MEFs. Human SRC residue numbering is used hereafter for clarity and consistency. In response to PV, reintroduction of SRC^WT^ increased global pTyr levels and phosphorylation of the SRC substrate cortactin, however, this was impaired for SRC^C188A^ and SRC^C280A^ indicating a role for these cysteines in driving signaling in response to PV (**Figs. 2J and 2K**). Importantly, PTP catalytic cysteine hyperoxidation detected by oxPTP was comparable in all cell lines. Together, our data suggest that oxidation of Cys 188 and 280 by PV are important for activating SRC and driving cellular increases in pTyr-containing substrates.

### PV triggers conformational changes that relieve SRC autoinhibition

We next performed a mechanistic investigation into SRC activation by oxidation. Previous molecular dynamics simulation suggested that cysteine oxidation plays a role in relieving autoinhibition (*21*), a key step in SRC activation. SRC autoinhibition is mediated by intramolecular contacts between SH2 and SH3 domains and the kinase domain (*35, 36*), with extensive conformational changes occurring during activation. To evaluate whether oxidation promotes conformational changes in SRC, we initially used a thermal shift assay using affinity purified proteins. ATP and Dasatinib stabilized full-length recombinant SRC, whereas PV and H_2_O_2_ led to destabilization, indicating oxidation-induced conformational changes (**Fig. S4A**).

To obtain a detailed molecular understanding of conformational changes caused by PV, we used hydrogen-deuterium exchange MS (HDX-MS) on full-length human SRC. The relative deuterium uptake differences were mapped onto the ‘closed’ (2SRC (*37*)) and ‘open’ (1Y57 (*38*)) SRC conformations (**Figs. 3A-3B**). At 0.5 mM (100x molar ratio of PV to SRC), PV increased solvent exposure of the SH2-kinase domain linker, which spans the polyproline-II helix. This helix engages the SH3 domain in the autoinhibited conformation and is released upon activation (**Figs. 3A *right,* and S4B**) (*35, 36*). In the SH2 domain, PV induced exposure of the αA helix while protecting the βB-βC loop, indicating redox-sensitive conformational changes near Cys 188. This suggests that 0.5 mM PV drives two key structural alterations involved in SRC activation.

**Fig. 3.**
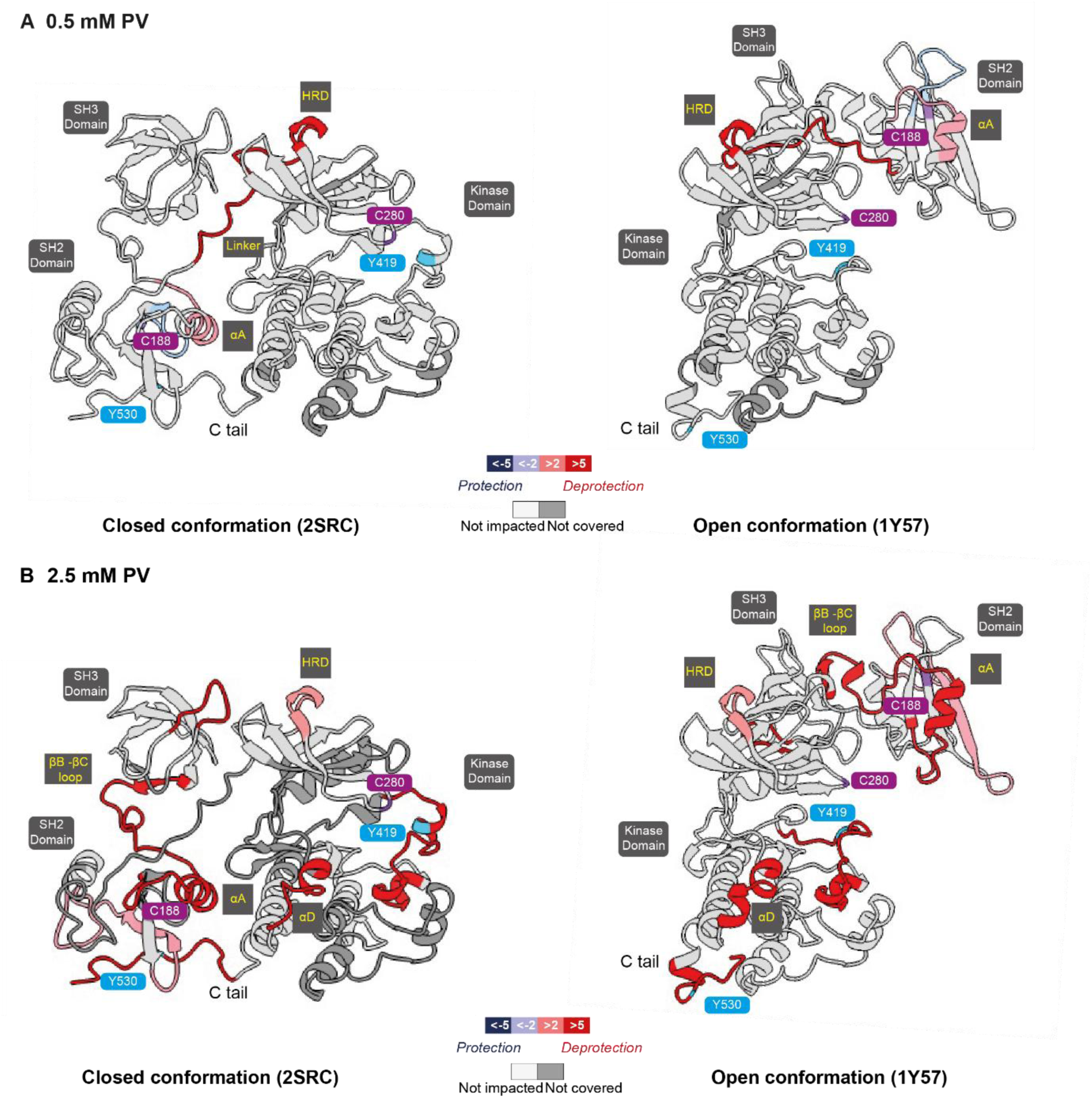
PV induces specific conformational changes in SRC. **(A, B**) HDX-MS analysis of pervanadate treated SRC. Relative fractional uptake (RFU) differences between 5 µM SRC and SRC incubated with 0.5 mM (**A**) and 2.5 mM (**B**) PV were mapped on the closed (Left) and the open conformations (Right) of SRC (PDB: 2SRC and 1Y57, respectively). Regions showing significant protection and becoming less solvent exposed are colored in blue and regions showing significant deprotection and becoming more solvent exposed are presented in red. Regions where no significant differences were observed are shown in white, while grey denotes parts of SRC for which no peptides were observed. Key cysteines are highlighted in purple and key phosphorylation sites are highlighted in light blue. The overall coverages were 80% and 60% for the 0.5 mM and 2.5 mM PV HDX-MS experiments, respectively. See also Fig. S4.

At 2.5 mM (500x molar ratio of PV to SRC), PV induced extensive solvent exposure across SRC regions associated with conformational transition from inactive to active kinase (**Fig. 3B**). The RT loop and the βE strand within the SH3 domain showed increased exchange, indicating release from the polyproline-II linker (*39*), a more pronounced effect than with 0.5 mM PV (**Fig. 3A**). Changes in the SH2 domain were also more prominent at this higher PV concentration (**Figs. 3B and S4C**). In the linker between the SH3 and kinase domains, a region spanning Trp 263 was exposed by PV. This key conserved residue is involved in forming a new β turn during the transition from closed to open conformations (*40*). The activation loop, including Tyr 419, was also more exposed aligning with previous MD simulations predicting Cys 280 oxidation-induced unfolding of this loop (*21*). The catalytic loop HRD motif and the αEF helix, which stabilize the activation segment in an active state, were also more exposed upon PV treatment (*37, 38, 41*) (**Fig. 3B, *right***). The C-terminal tail, including Tyr 530, was more accessible along with the αD helix (**Fig. 3B, *right***). Overall, PV induced conformational changes that increased flexibility and solvent accessibility, explaining the oxidation-induced destabilization observed by thermal shift (**Fig. S4A**). These effects were increased at 2.5 mM PV, consistent with the dose-dependent effects observed in kinase assays (**Fig. 2A**). Together, these results support a model in which PV drives the relief of SRC autoinhibition via specific conformational alterations across the polypeptide. Furthermore, at lower levels of oxidation we observed conformational changes largely in the vicinity of Cys 188, aligning with its greater sensitivity to oxidation (**Figs. 2E and 2F**), with effects mediated by Cys 280 requiring increased PV levels. This supports a hierarchy of cysteine PV sensitivity within SRC.

### Cys 188 oxidation impacts SRC-SH2 affinity for phosphopeptides

We next focused on how the conformational changes within the SRC SH2 domain revealed by our HDX-MS data may contribute to relief of autoinhibition (**Fig. 4A**). Cys 188 resides within the pTyr binding pocket and is ideally placed to mediate SRC interactions with phosphopeptides (**Figs. 4A-4B**). We used AlphaFold3 to explore the binding of SRC-SH2 domain with the phosphorylated SRC C-terminal peptide, which interact in the autoinhibited state (*42*). Cys 188 is predicted to participate in a hydrogen bonding network that engages the phosphate group of the bound peptide (**Figs. 4C, *Top* and S5A**). To mimic the effects of oxidation, we modelled Cys 188 with an S-nitroso modification in AlphaFold3 (**Figs. 4C, *middle* and S5B**). We infer that Cys oxidation causes extensive remodelling of the hydrogen-bonding network, reducing the overall number of contacts with the phosphate group. In contrast, modelling C188A predicts hydrogen bonding network adjustments that incorporate new interactions, including with Thr 182 (**Figs. 4C, *bottom* and S5C**), which might impact phosphopeptide binding.

**Fig. 4.**
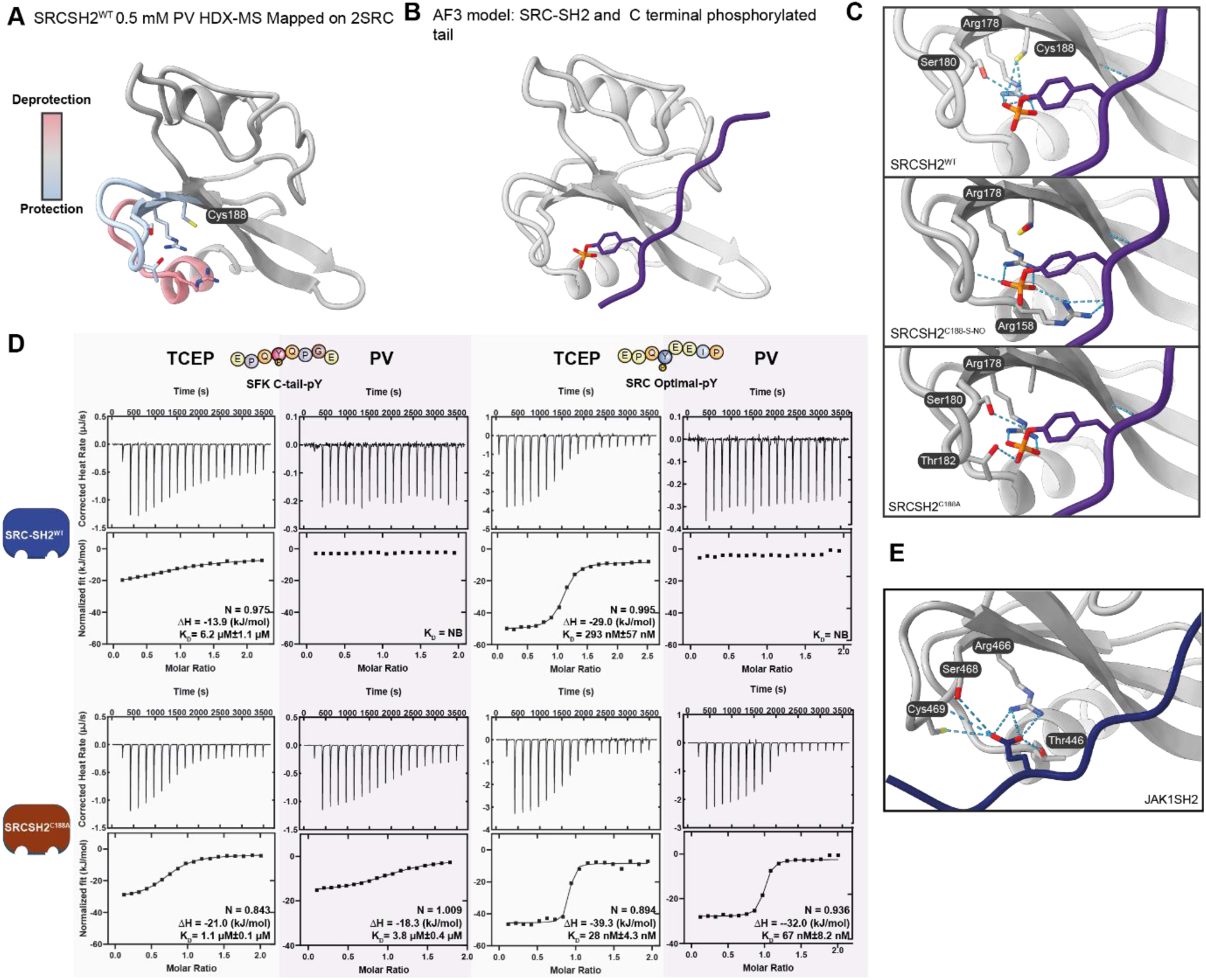
Cys 188 oxidation or mutation impacts SRC-SH2 phosphopeptide binding. **(A)** SRC-SH2 with highlighted regions protected (blue) and deprotected (red) from deuterium exchange upon 0.5 mM PV treatment (PDB:2SRC) based on data from Fig. 3. (**B**) Alphafold3 (*42*) model of SRC SH2^WT^ bound to the phosphorylated SRC C-terminal peptide: FTSTEPQpYQPGENL. ipTM = 0.73. (**C**) Residues predicted to form hydrogen bonds (blue dashed lines) based on AF3 models. Top: As in B. Middle: AF3 model with SRC s-nitroso cysteine 188 and C terminal peptide. ipTM = 0.75. Bottom: AF3 model with C188A and C terminal peptide. ipTM = 0.72. (**D**) Isothermal titration calorimetry (ITC) titration curves of the interaction between SRC-SH2 and indicated phosphopeptides. The SH2 domain was pretreated with 1 mM TCEP or 0.1 mM PV prior to the assay. Upper: baseline-corrected differential power plotted over time. Lower: normalized binding curve showing the integrated change in enthalpy against the molar ratio. Stoichiometry (N), mean enthalpy change (ΔH) and mean K_D_ ±SD shown based on n =3 assays. (**E**) AF3 model of hydrogen bonding network between residues within the JAK1 SH2 domain engaging glutamate on IFNR1. ipTM = 0.69. See also Fig. S5.

We next used isothermal titration calorimetry (ITC) to quantify effects of PV on phosphopeptide interaction dynamics using WT and C188A SRC-SH2 domains (**Fig. 4D**). As predicted by our AlphaFold3 analysis (**Fig. 4C, *bottom***), under reducing conditions (1 mM TCEP), SRC-SH2^C188A^ showed a 5-fold increase in affinity for the SRC C-terminal phosphopeptide (spanning pY530) compared to SH2^WT^ (**Fig. 4D**). Strikingly, oxidation of SH2^WT^ with 0.1 mM PV abolished phosphopeptide binding, consistent with uncoupling of the inhibitory complex. In marked contrast, there was still detectable binding to SH2^C188A^, albeit with ∼3.5-fold lower affinity than under reducing conditions. Next, we examined interactions with an optimised SRC binding phosphopeptide (EPQ**pY**EEIP) (*43*). We detected a ∼10-fold increase in affinity with SH2^C188A^ compared to SH2^WT^, consistent with a previous study (*44*) (**Fig. 4D, *lower***). Strikingly, PV treatment eliminated phosphopeptide binding by SH2^WT^, but not SH2^C188A^, which showed a ∼3-fold decrease in affinity compared to TCEP-treated samples (**Fig. 4D**). Both Cys 241 and Cys 248 are present within the SH2 domain and may contribute to this residual PV sensing. Nevertheless, our results support Cys 188 in mediating SRC-SH2 phosphopeptide binding and sensitivity to PV. These findings provide a plausible mechanistic explanation for the impact of Cys 188 oxidation on the activity of SRC in cells.

Finally, we asked whether other human SH2 domains possess putative redox sensitive cysteines that could alter phosphopeptide interactions. Interestingly, SH2D1A, SH2D1B and SH2D5 all possess a cysteine analogous to SRC Cys 188 (**Fig. S5D**). Furthermore, a mutation affecting this cysteine in the SH2D1A domain is linked to lymphoproliferative disease (*45*). Several signaling proteins also possess SH2 domains with cysteine residues within their pTyr binding pockets (*46*) (**Fig. S5E**), including the PTKs Zap70 and Syk, which exhibit reduced phosphopeptide binding following oxidation by H_2_O_2_ (*47*). We anticipate that cysteine residues (*46*) within the CLNK and JAK1-SH2 domains would also regulate signaling in a redox-sensitive manner (**Figs. 4E and S5E**). Thus, redox-regulated SH2 domain-phosphopeptide interactions are important beyond SRC.

### Mutational relief of SRC autoinhibition bypasses the functions of Cys 188 and Cys 280 in cells

Mutating SRC Tyr 530 (Y530F) favours its open, active conformation (*48*). We hypothesized that this mutation would render SRC less sensitive to the effects of cysteine mutation and activation by oxidation. To test this, we introduced SRC^Y530F^ and dual mutant variants SRC^Y530F;C188A^ and SRC^Y530F;C280A^ into SYF KO MEFs. As expected, basal pY419 and global pTyr signal in cells expressing SRC^Y530F^ was increased compared to those expressing SRC^WT^ (**Figs. 5A and S6A**). SRC^C188A^ and SRC^C280A^ showed a striking increase in pY530 and reciprocally diminished pY419, indicating increased autoinhibition. In contrast, introducing the Y530F mutation overcame the inhibitory effects of C188A and C280A in unstimulated cells, as determined by pY419 levels (**Figs. 5A and S6A**). PV stimulated global pTyr equivalently for all SRC^Y530F^ variants independently of Cys 188 or Cys 280 mutations. In contrast, PV-induced pTyr levels were lower for SRC^WT^ cysteine mutations. We also noted that SRC pY530, a reported substrate of numerous PTPs (*49-51*), was unchanged by PV treatment (**Fig. 5A**). These results support that Cys 188 and Cys 280 regulate SRC activity at the level of autoinhibition in cells. Aligning with this, SRC Cys 188 and Cys 280 have been implicated in transducing signals downstream of ATP (*21*) and IL33 (*52*) stimulations.

**Fig. 5.**
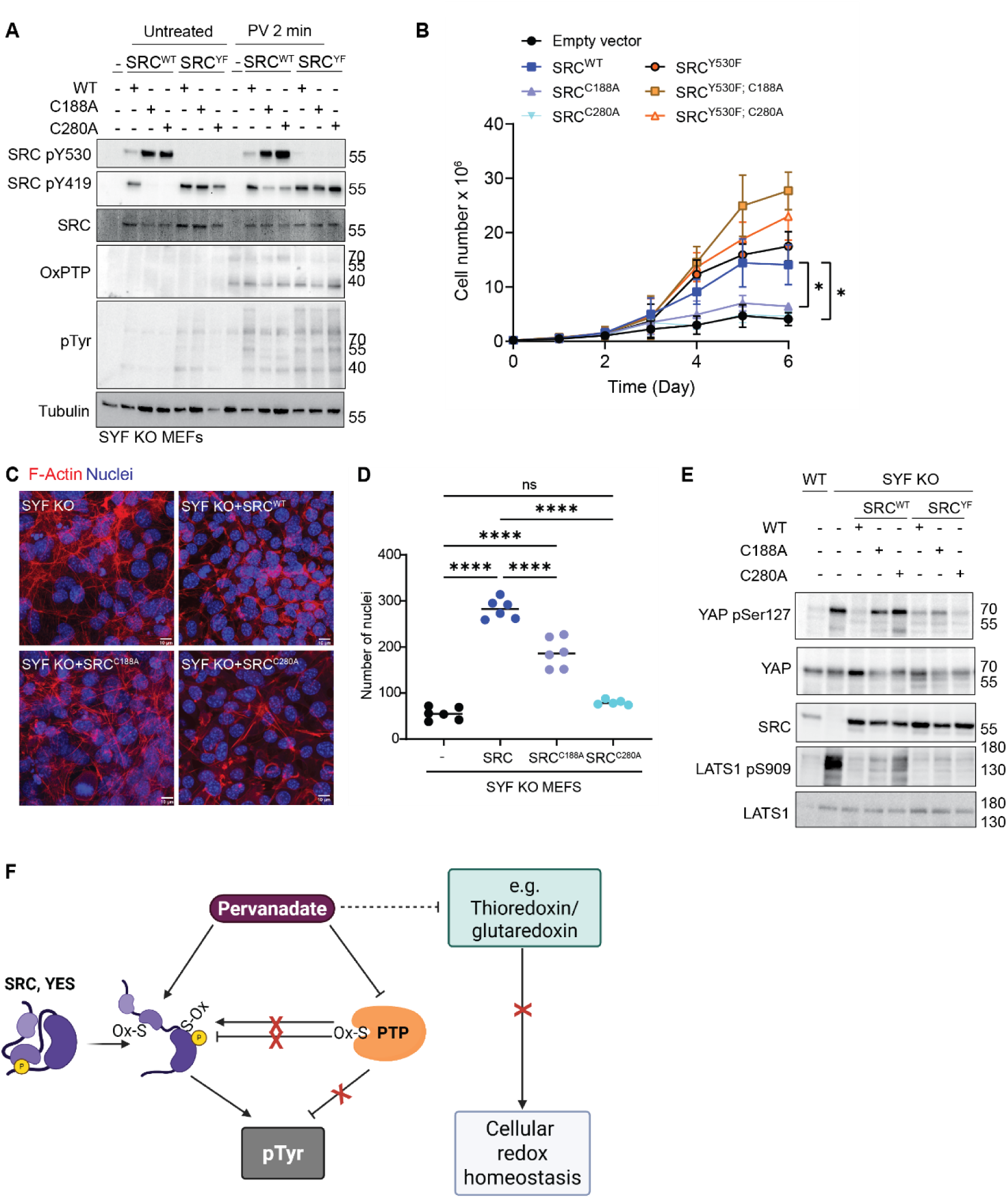
SRC cysteines are required to overcome contact inhibition. (**A**) SYF KO MEFs were retrovirally transduced with specified SRC expression constructs and treated with 100 µM PV as shown, before lysis and immunoblotting with indicated antibodies. (**B**) Cell numbers were determined over time in culture for SYF KO MEFs, retrovirally transduced with specified SRC expression constructs (as in A). (**C**) Confocal microscopy images of SYF KO MEFs complemented with the indicated constructs grown to post-confluence, fixed, permeabilized and stained with the DAPI (nuclei; blue) and Phalloidin (actin; red). (**D**) Quantification of nuclei taken from 5 or 6 fields of view per cell line. **** p.Adj < 0.001. Ordinary one-way ANOVA with Tukey’s correction for multiple comparisons. (**E**) SYF KO MEFs expressing the indicated constructs were grown to post-confluence, lysed and immunoblotted with the indicated antibodies. Representative blot from n=3 experiments. (**F**) In summary, Pervanadate induces numerous cellular changes. Increases in pTyr can be attributed to the hyperoxidation of PTP catalytic cysteines as well as direct oxidation of specific cysteines on SRC and YES leading to relief of autoinhibition and activation. Furthermore, PV induces rapid oxidation of HyPer7, a reporter for both H2O2 and the activity of cellular antioxidant systems such as thioredoxin and glutaredoxin. PV therefore disrupts redox homeostasis in addition to increasing phosphotyrosine levels. See also Fig. S6.

### SRC Cys 188 and Cys 280 are required for overcoming cell contact inhibition

Reintroducing SRC^WT^ into SYF KO MEFs stimulated proliferation (**Fig. 5B**). This effect was not recapitulated by expression of SRC^C188A^ or SRC^C280A^, reflecting the reduced basal cellular activity of these mutants (**Fig. 5B**). In contrast, constitutively active SRC^Y530F^ increased cell growth regardless of C188A and C280A mutations. Notably, SRC^Y530F; C188A^ cells exhibited the highest growth rate (**Fig. 5B**), potentially reflecting a negative impact of Cys 188 oxidation, which would impair pY530 binding but would subsequently also limit binding to and recruitment of other phosphoproteins (**Fig. 4**). We observed that differences in cell number were largely driven by post-confluent growth. Indeed, SRC^WT^ and SRC^Y530F^ expressing cells did not undergo contact inhibition, whereas SYF KO, SRC^C188A^ and SRC^C280A^ cell lines did (**Fig. 5B and movies S3-S6**). Following growth to post-confluence, SRC^WT^ expressing cultures had significantly more nuclei supporting increased cell proliferation compared to either the SYF KO or SRC^C188A^ and SRC^C280A^ (**Figs. 5C-5D**). Together, these findings demonstrate that SRC Cys 188 and Cys 280 are required to drive cell overgrowth.

The Hippo pathway is a conserved regulator of contact inhibition of cell growth (*53*). We next tested whether SRC-dependent cellular overgrowth phenotypes were associated with YAP activation (*53*). Compared to WT, SYF KO MEFs displayed increased YAP phosphorylation at Ser 127 (**Fig. 5E**), a site associated with its cytoplasmic sequestration and inactivation. This trend was recapitulated by phosphorylation of the upstream Hippo pathway kinase LATS1 (**Figs. 5E and S6B**). Consistently, expression of SRC^WT^ was sufficient to suppress YAP phosphorylation, whereas SRC^C188A^ and SRC^C280A^ resembled SYF KO cells, with elevated levels of YAP phosphorylation, despite comparable SRC expression levels (**Fig. 5E**). In conclusion, SRC redox-sensitive cysteines are required for YAP activation, correlating with the ability of cells to overcome contact inhibition of proliferation.

## Discussion

PV is a commonly used PTP inhibitor that has shaped our understanding of pTyr-based regulation and the study of PTP-substrate interactions (*54, 55*). Our work demonstrates that PV can disrupt cellular redox homeostasis, leading to oxidation of HyPer7 and PRDX1, but also shows selectivity, directly targeting cysteines within positively charged protein surfaces. Using a series of mutational approaches, enzyme assays and cellular studies, we find that the potent effects of PV on global pTyr are best explained by the combined inhibition of PTPs and the direct oxidation and activation of SFKs, and perhaps other PTKs, such as Eph receptors (*56*). This finding should prompt a re-evaluation of the use and interpretation of experimental data generated using PV treatments.

Based on the oxidation of both HyPer7 and PRDX1, PV must interfere with antioxidant pathways, particularly the redoxins that can directly or indirectly reduce both proteins. One possibility is that it reacts with and depletes reduced glutathione (GSH) or NADPH. PV probably evades H_2_O_2_-reactive PRDX, enabling it to target additional cysteine thiols similarly to what has been observed for *cis*Trp-OOH (*57*). This would be consistent with rapid PTP hyperoxidation. Over time, PV induces PRDX1 tyrosine phosphorylation, potentially enabling endogenous H_2_O_2_ to oxidise cysteine thiols (*32*). Interestingly, the SRC cysteine residues studied here showed a distinct pattern of sensitivity to H_2_O_2_ than we observe for PV (*21*). Therefore, we propose that PV preferentially targets PTP active sites, as well as other phosphate-binding pockets, including the SRC kinase active site (Cys 280) and SH2 domain (Cys 188), correlating with protein regions with high positive charge (**Fig. 2G**). In contrast, mercuric chloride activates SRC independently of autoinhibition via Cys 486 and 501, which are surrounded by negative charge, particularly in the open conformation (*58*). Therefore, SRC responds differentially to distinct oxidants.

Cys 188 is located at an equivalent position to Thr 42 in the N-terminal SHP2 SH2 domain, which is mutated in disease and alters phosphopeptide specificity (*59*). Likewise, SRC-SH2^C188A^ altered affinity for phosphopeptides, but also impacted redox sensitivity. Interestingly, Cys 188 reportedly forms a disulfide bridge with a cysteine residue of its substrate cortactin (*60*), which could explain the greater influence of mutating this cysteine in cells than *in vitro*. Cys 188 oxidation is likely to play two roles: firstly, decreasing affinity for phosphopeptides and promoting relief of autoinhibition. and secondly, mediating disulfide bridge formation with interaction partners, an intriguing area for further investigation. Amongst SFK cysteine residues, Cys 188 exhibits the highest level of reversible oxidation in mouse tissues, supporting the physiological relevance of our findings (*20, 61*). We further identified other SH2 domain pTyr pocket cysteine residues. Zap70 possesses a redox-sensitive pTyr binding pocket in its N terminal SH2 domain, but has a distinct mode of autoinhibition compared to SRC (*62*). Thus, oxidation of this cysteine may negatively impact ZAP70 recruitment to phosphorylated immunoreceptor tyrosine-based activation motifs (ITAMs) (*47*) but may also mediate intermolecular disulfide interactions. Interestingly, CSK, which promotes SRC Tyr 530 phosphorylation and subsequent inhibition, is regulated *in vitro* via an intramolecular disulfide bond involving its SH2 domain and could also contribute to redox and pTyr signalling integration (*63*). Thus, redox regulation of SH2 domains is a wider area for future investigation.

SRC^C188A^ and SRC^C280A^ expressing cells showed a striking increase in pY530, which differs from previous reports (*21*). One explanation for this difference is that we ‘rescued’ SRC-deficient cells, whereas other studies were performed in the presence of endogenous SRC (*21*). Furthermore, we have employed untagged SRC wherever possible, since both the N and C termini of SRC contain important regulatory elements. For example, a C terminal affinity tag could interfere with autoinhibition, whilst N-terminal tags would block N-myristoylation, which is important for localisation (*64*). This striking increase in pY530 (**Fig. 5A**) may be consistent with the preservation of an autoinhibited state and protection from dephosphorylation by a stabilized SH2 domain interaction. Alternatively, it could represent an increase in CSK activity, although this has not been mechanistically explored in this study.

In *Drosophila*, Src42A Cys 471, equivalent to human SRC Cys 490, is implicated in wound healing responses (*65*), and may form an intramolecular disulphide with Cys 248 (*19*). Neither Cys 188 nor Cys 280 are conserved in *Drosophila* Src (42A or 64B), but are conserved in zebrafish and chicken, indicating multiple layers of SRC redox regulation acquired during evolution. SRC can be S-glutathionylated at Cys 400 leading to inhibition of autophosphorylation, linking its cellular regulation to this pathway (*66*). How SRC cysteines undergo oxidation and reduction *in vivo* remains an important question, particularly given the relatively high oxidant concentrations required to elicit changes in activity *in vitro*.

SRC is a proto-oncogene yet rarely mutated in human cancers (*67*). Cys 188 and Cys 280 mutations prevent SRC mediated cell overgrowth, strongly suggesting that the oxidative environment typical of some cancers (*68*) could drive relief of SRC autoinhibition, phenocopying Y530F or C-terminal truncations (*67*). Our findings with the conserved Cys 280 residue also have implications for YES1, for which we also demonstrate oxidative activation by PV (**Fig. S3G**). Interestingly, the equivalent cysteine in YES1 (Cys 287) has previously been targeted with a covalent inhibitor (*69*). Our finding that non-catalytic redox-sensing Cys residues in PTKs is a dominant signaling mechanism potentially increases their attractiveness as therapeutic targets, especially if greater potential for selectivity than active site inhibitors can be achieved (*70*).

## Materials and methods

### Plasmids and constructs

Amino acid (aa) numbering is based on the following sequences: Human PTPRK; UniProt ID: Q15262-3, Human SRC; Uniprot ID: P12931-1, Chicken Src Uniprot ID: P00523. Generation of bacterially expressed (aa 864–1446) and mammalian overexpression PTPRK intracellular domain (aa 776-1446) constructs were previously described (*73, 74*). Full length human SRC containing a 3C-protease cleavable, N-terminal, tandem Strep-tag II was cloned into pcDNA3. Human rhinovirus 14 3C protease plasmid (pET-NT*-HRV3CP) was a gift from Gottfried Otting (Addgene # 162795).

The retroviral pLNCX chicken Src Y527F plasmid was a gift from Joan Brugge (Addgene #13660). HyPer7 (*30*) (a gift from Vsevolod Belousov (Addgene #136464)) was fused in-frame downstream of the SRC myristylation sequence to make the ‘PM-HyPer7’ biosensor and subcloned into pLJM1 (gift from Joshua Mendell (Addgene #91980)). N-terminally His-tagged HyPer7 was cloned into the pQE30 vector. The cDNA encoding the human SRC SH2 domain (Gln147-Lys252) was subcloned into the pETM11 vector (a gift from Elisa Izaurralde (Addgene #145950)). The QuikChange II mutagenesis kit (Agilent) was used to construct the mutant SRC SH2 domains.

All other point mutations were introduced by polymerase chain reaction (PCR) using either Q5 High-Fidelity DNA (New England Biolabs, UK), CloneAmp (Takara) or Phusion Hot Start II DNA (Thermo Fisher Scientific) polymerases as per manufacturer’s protocol.

### Cell culture and treatments

HEK293T, U2OS and SYF KO MEFs (CRL-2469) cells were purchased from ATCC. WT MEFs were generated in-house, described below. SRC WT and KO HAP1 cells were purchased from Horizon (HZGHC000348c023). Plat-E cells were donated by P. Hawkins and L. Stephens (Babraham Institute) and were purchased from Cell Biolabs Inc (RV-101). Cells were maintained in 75 cm^2^ vented tissue culture flasks in a 37°C humidified 5% CO_2_ ventilator and passaged using trypsin-EDTA solution (Sigma-Aldrich), prior to reaching confluence every 2-4 days. HEK293T and MEF cells were grown in DMEM containing 10% (v/v) fetal bovine serum (FBS; Sigma-Aldrich), 2 mM L-glutamine (Sigma-Aldrich). U2OS cells were grown in McCoy’s 5A medium modified with 10 % (v/v) FBS (Sigma-Aldrich). HAP1 cells were cultured in IMDM containing 10% (v/v) fetal bovine serum (FBS; Sigma-Aldrich, with Plat-E cells cultured in DMEM (high glucose with L-glutamine and pyruvate supplemented) containing 10 % heat-inactivated FBS (Sigma-Aldrich) with additional blasticidin (10 mg/ml) and puromycin (1 mg/ml) when in culture for more than 2 passages. All cell lines are routinely mycoplasma tested using MycoProbe Mycoplasma Detection Kit (R&D system) or MycoAlertTM (Lonza Bioscience).

### MEF line generation

WT (Bl6/N) MEFs were generated in-house. Following dissection, E13-14 embryos were placed into PBS on ice. Placental and maternal tissues and head were removed prior to tryspinization and dissociation with a razor blade. DNAseI was then added followed by incubation at 37°C for 10 min. DMEM with 10 % FBS was added to quench trypsin. Cells were pipetted through a 40 µm filter and pelleted at 140 x g for 5 min. Finally, cells were resuspended in DMEM with 10% FBS, 2 mM L-glutamine and pen-strep then seeded for culturing. Cells were cryopreserved in liquid N2 in 50% FBS:10% DMSO:40% DMEM for future use.

### Small molecules and growth factors

The following inhibitors were purchased from Sigma: Dasatinib (SML2589-50MG), focal adhesion kinase inhibitor 1 (324877-5MG), Gefitinib (SML1657-10MG), Ibrutinib (PZ0242-5MG), Imatinib (CDS022173-25MG), Lapatinib (SML2259-10MG), Nilotinib (SML3799-10MG), PP2 inhibitor (529573-1MG), Saracatinib (SML3195-5MG) and SU6656 (S9692-5mg). Inhibitors were re-suspended in DMSO. The following growth factors were purchased from Qkine: PDGF (Qk044), IGF1 (Qk047), FGF2 (Qk025), TGFβ2 (Qk072). Insulin was purchased from Scientific Laboratory supplies (I5500) and EGF was purchased from Peprotech (AF-100-15).

### Antibodies and dyes

Antibodies against the following target proteins or epitopes were used in the study: Phosphotyrosine (Cell Signaling Technology (CST) #8954S), α Tubulin (Sigma #T6199), Flag (CST #8146S), Pan-SFK (CST #2108), SRC family pY416 (#CST D49G4), SRC pY530 (CST #2105), SRC-specific (CST #2123 & #2120), oxPTP (R&D Systems # MAB2844), His (CST #12698S), ABL1 (c-Abl) (CST #2862), ABL1 (c-Abl) pY412 (CST #2865S), Cortactin (CST #3503), Cortactin pY421 (CST #4569), LATS1 (CST #3477), LATS1 pY909 (CST#9157), YAP (CST #4912), YAP pS127 (CST #13008), AKT (CST #9272), AKT pS473 (CST #4060), ERK pT202/pY204 (CST #4370), ERK (CST #4696), EGFR (CST #2232S), EGFR pY1068 (CST #3777S), SHP2 (CST #3397), PRDX1 pY194 (CST #14041), PRDX1 (CST #8499). HRP-conjugated secondary antibodies were purchased from Jackson Immunoresearch(#715-035-150 & #711-035-152). Hoechst and BODIPY Phalloidin 558/568 (B3475) was purchased from Thermo Fisher Scientific.

### Pervanadate stock generation

Pervanadate was generated as previously described (*6*). To avoid decomposition, it was prepared immediately prior to use. In a 1.5 ml tube, 100 μl of activated pH 10 100 mM sodium orthovanadate (Alfa Aesar 15403009) was combined with 103 μl 0.49 M H_2_O_2_ in 20 mM HEPES, pH 7.3, mixed by gentle inversion and incubated at RT for 5 min. Excess H_2_O_2_ was quenched by addition of 23 μl of 0.5 mg/ml catalase in 50 mM potassium phosphate, followed by mixing by gentle inversion, yielding a 44.2 mM pervanadate stock solution for immediate use.

### Recombinant proteins and synthesized peptides

Peptides were obtained at 95% purity from Biosynth. The SRC peptide substrate 5-FAM-EGIYGVLFKKK-CONH2 was synthesized by Genscript. Recombinant N-terminal His6-tagged SRC (1-536) kinase fusion protein (DU19041) and N-terminal His6-tagged YES1 (1-543) (DU5884) was purified from Sf21 cells was purchased from Medical Research Council Protein Phosphorylation and Ubiquitylation Unit (PPUU) reagents and services (University of Dundee). N-terminally His-tagged Hyper7 (*30*), PTPRK ICD (*73*), SRC SH2 domain and His6-tagged 3C protease (*75*) were purified using standard affinity and size exclusion chromatography methods and stored −80°C as follows*: M15 Escherichia coli* cells were transformed with N-terminally His-tagged Hyper7 construct and cultured in LB Broth at 37°C to OD600nm ∼0.5-0.8 and induced with 400 µM isopropyl-thio-β-D-galactopyranoside (IPTG Thermo Fisher Scientific, dioxane-free) for 14 h at 16°C. *Escherichia coli* BL21(DE3) Rosetta cells transformed with SRC SH2 construct or the other aforementioned constructs were cultured at 3<u>0</u>7°C/220 rpm in LB broth containing 34 μg/ml chloramphenicol until the OD600 nm reached 0.5–0.8 or cultured at 30°C/220 rpm in 1 L 2XTY medium containing 50 μg/ml carbenicillin and 34 μg/ml chloramphenicol until the OD600 reached 0.6–0.7 respectively. Cultures for SRC SH2 were transferred to 16°C/220 rpm and induced with 100 µM IPTG (Thermo Fisher Scientific, dioxane-free) for 14 h. All other constructs were transferred to 20°C/220 rpm and allowed to equilibrate, prior to the addition of 1 mM IPTG (Generon) for 20h. PTPRK cultures were also supplemented with 200 μM of D-biotin (Sigma-Aldrich) upon IPTG addition. Cells were harvested by centrifugation at 4000 x g for 20-30 mins and bacterial pellets were stored at – 20°C until required.

For Hyper7, pellets were resuspended in lysis buffer (40 mM HEPES, pH 7.4, 400 mM NaCl, 10% (v/v) glycerol, 1 mM benzamidine, 1 mg/mL Leupeptin, 0.1 mg/mL 4-(2-aminoethyl) benzenesulfonyl fluoride hydrochloride (AESBF), 100 mg/mL DNaseI, 20 mM MgCl2) or oxidizing or reducing lysis buffer for SRC SH2 domains (20 mM HEPES (pH 7.5), 100 mM NaCl, and 0.1 mM pervanadate; reducing SH2 buffer: 20 mM HEPES (pH 7.5), 100 mM NaCl, and 1 mM TCEP). Once re-suspended in lysis buffer, samples were lysed by sonication and clarified by centrifugation at 39,000 x g for 30 min at 4°C. For PTPRK and 3C protease proteins, cells were lysed in purification buffer (50 mM HEPES pH 7.5, 500 mM NaCl, 5% (v/v) glycerol and 0.5 mM tris (2-carboxyethyl)phosphine (TCEP)), containing EDTA-free protease inhibitor tablets (Roche) followed by cell disruption using a Constant Systems cell disruptor and then lysate clarification via centrifugation at 40000 x g for 30 min at 4°C. His-HyPer7 were purified by affinity chromatography (Ni2+-Sepharose (GE Healthcare) and imidazole-based elution) and size exclusion chromatography using a HiLoad 16/600 Superdex 200 column (GE Healthcare) equilibrated in 50 mM Tris-HCl (pH 7.4), 100 mM NaCl, and 10% (v/v) glycerol. SH2 domains were purified with pre-equilibrated nickel column (His-Trap HP, GE Healthcare) and the 6x His-tag was removed by digestion with home-made tobacco etch virus protease at 4°C for 12 h accompanied with oxidizing buffer and reducing buffer.

SH2 domains were further purified on a Superdex 75 10/300 GL size exclusion column (GE Healthcare) with oxidizing buffer and reducing buffer. For PTPRK and 3C protease, the supernatant was removed and incubated with 0.5 ml of Ni-NTA agarose (Qiagen) for 1 h at 4°C. Ni-NTA Agarose was then pelleted via centrifugation at 500 x g for 5 min at 4°C and packed into a gravity flow column. Ni-NTA agarose was then washed with 10 volumes of purification buffer containing 5 mM imidazole, followed by 20 volumes of purification buffer containing 20 mM imidazole; prior to elution in purification buffer containing 250 mM imidazole. The eluted protein was then subjected to size exclusion chromatography (SEC) using a Superdex 200 16/600 column (GE Healthcare Life Sciences, Thermo Fisher Scientific). Columns were equilibrated in SEC buffer (50 mM HEPES pH 7.5 (50 mM Tris pH 7.4 for SH2 domains), 150 mM NaCl, 5% (v/v) glycerol, 5 mM DTT). Protein was concentrated to 2–10 mg/ml using an Ultracel-10K regenerated cellulose centrifugal filter (Merck Millipore). Proteins were either used fresh or snap-frozen for storage at -80°C until required.

### Generation of stable cell lines

SRC-rescued SYF KO MEF cell lines were generated using the retroviral expression system. Retrovirus was generated through the reverse transfection of Plat-E cells (12 x 10^6^) in 15 cm^2^ dishes with 16.8 µg pLNCX construct DNA into 2 ml OPTIMEM (Gibco) with 50.4 µl Lipofectamine 2000 transfection reagent (Invitrogen) following the manufacturer’s instructions. Media was replaced with 9 ml complete MEF media per 15 cm^2^ dish 20-24 hours post-transfection. 48-72 h post-transfection, retrovirus was harvested from Plat-E cell media by centrifugation for 5 min at 800 x g (RT) and filtered through a 0.2 μm syringe filter (Sartorius) before either freezing at -80°C or adding directly onto MEF SYF KO cells after dilution 1:1 into MEF culture media, in 6-well dishes. Media was replaced 24 h after adding viral supernatant, followed by at least two passages before experimental analysis.

For lentiviral generation, 1.5×10^7^ HEK293T cells were seeded in 12 ml of complete growth medium per 15 cm^2^ dish (two dishes per lentivirus) and incubated for 24 h at 37°C with 5% CO_2_. Each 15 cm^2^ dish was then transfected with 6 μg of pLJM.PM-HyPer7, 12 μg psPAX2 packing plasmid (Addgene #12260, deposited by Didier Trono) and 3 μg pMD2.G envelope plasmid (Addgene #12259, deposited by Didier Trono) using the GeneJuice transfection reagent (Merck Millipore, UK) as per manufacturer’s instructions. After 24 h, the medium was then replaced with 16 ml complete growth medium. At 48–72 h post-transfection, culture medium was collected and filtered through a 0.2 μm syringe filter (Sartorius). Viral particles were concentrated using Lenti-X concentrator as per manufacturer’s instructions (Takara Bio). Lentivirus was centrifuged at 1500 x g for 45 min at 4°C, resuspended with complete DMEM media and aliquoted and stored at −80°C until required. 2×10^5^ U2OS cells were seeded per well of a six-well plate in 900 μl of growth medium, prior to the drop-wise addition of 100 μl concentrated lentivirus. After 30 min at room temperature, the cells were returned to the incubator. 72 h later, cells were reseeded in 1 μg/ml puromycin (Thermo Fisher Scientific) selection medium. Cells were subsequently maintained in media supplemented with 1 μg/ml puromycin.

### Lipid-based transfection of siRNA pools

MEF SYF KO cells were reverse transfected with siRNAs (ON-TARGETplus SMARTpool, Dharmacon Horizon Discovery) using lipofectamine RNAiMAX as per the manufacturer’s instructions (Thermo Fisher Scientific 13778030). Briefly, for each well of a 6-well plate, 6 µl RNAiMAX was used to transfect 10 nM siRNA. After 24 hr, media was replaced with complete growth medium and left to recover for another 24 hr prior to cell treatments and processing for analysis.

### pNPP phosphatase activity assay

Recombinant PTP domains were made up to 500 μl in assay buffer (50 mM Tris–HCl, pH 7.4, 150 mM NaCl, 5% [v/v] glycerol, 5 mM DTT) at 1 μM. PTPRK intracellular domain and 20 mM pNPP substrate (NEB P0757S) (in assay buffer) were equilibrated to 30°C for 15 min in an orbital shaking heat block at 500 RPM. To initiate reactions, 500 μl of 20 mM pNPP substrate was added to PTP containing tubes (0.5 μM PTP and 10 mM pNPP final concentrations). Reactions were carried out for 120 min at 30°C in an orbital shaking heat block at 500 RPM. At each time point (0, 2, 5, 10, 15, 30, 60, 90 and 120 min), 100 μl of the total reaction was transferred to a 96-well microplate well containing 50 μl 0.58 M NaOH, terminating the reaction. After the final time point, absorbance of each sample was measured at 405 nm in a Spectramax M5 plate reader (Molecular Devices). Product formation was calculated by interpolation of absorbance values using a 4-nitrophenol standard curve of known concentration.

### SDS-PAGE and immunoblotting

Protein or cell lysate samples were resuspended in an appropriate volume of 5X SDS-PAGE sample buffer (0.25 M Tris-HCl pH 6.8, 10% (w/v) SDS, 20% (v/v) glycerol, 0.1% (w/v) bromophenol blue, 10% (v/v) β-mercaptoethanol) and incubated at 92°C for 10 min. Samples were run on a 10% (v/v) SDS-polyacrylamide resolving gel with a 5% (v/v) SDS-PAGE stacking gel and subjected to electrophoresis at 125 V for 1.5 h in 25 mM Tris, 190 mM glycine, 0.1% (w/v) SDS. Proteins were transferred onto 0.2 µm reinforced nitrocellulose membranes (GE Healthcare) at 300 mA for 3 h at 4°C in 25 mM Tris, 190 mM glycine, 20% (v/v) methanol. Membranes were briefly rinsed in TBS-T (20 mM Tris pH 7.6, 137 mM NaCl, 0.1% (v/v) Tween-20) prior to incubation for 30 min in 5% (w/v) skimmed milk/TBS-T to block non-specific antibody binding. The blocking solution was removed and membranes rinsed in TBS-T prior to primary antibody incubation (overnight at 4°C). Membranes were then subjected to 3 × 10 min washes in TBS-T, prior to incubation with HRP-conjugated species-specific anti-IgG antibodies (1 h at RT). Membranes were then subjected to 3 × 10 min washes in TBS-T, prior to being incubated with combined EZ-ECL solution (Cytiva) and imaged using a G:BOX Chemi XX6 (Syngene).

### Peroxiredoxin disulphide shift

HEK293T cells were subjected to a pervanadate time course followed by incubation with PBS with or without 10 µg/ml catalase (SLS #C3155) and 100 mM NEM for 15 mins at room temperature. Catalase was added to the PBS 30 min prior to use with NEM added immediately prior to use. Samples were lysed in 2 % SDS lysis buffer with and without 10 mM NEM. Samples underwent immunoblotting on 12 % gels with either non-reducing or reducing conditions and probed with PRDX1.

### PTPRK Flag overexpression, immunoprecipitation and pNPP assay

Flag IP-phosphatase assays were performed as previously described (*74*). HEK293T (6 x10^6^) cells were plated in 10 cm^2^ dishes, 24 h prior to transfection. 8 µg expression constructs for 3x FLAG tagged ICDs were transfected using 24 µl GeneJuice (Sigma) in 1 ml OPTIMEM (Gibco) serum free media. Media was replaced 4 h post-transfection. For immunoprecipitation (IP), cells were transferred to ice 48 h post transfection and washed twice with ice-cold 1X phosphate buffered saline (PBS). Cells were lysed in ice-cold lysis buffer (50 mM Tris-HCl pH 7.4, 150 mM NaCl, 10 % [v/v] glycerol, 1 % [v/v] Triton X-100, 1 mM EDTA, 10 mM NaF, 1 mM PMSF, 1X-EDTA-free protease inhibitor) on ice for 30 mins with periodic agitation. Lysates were collected and cleared by centrifugation at 14000 x g for 15 min at 4 °C and supernatants were transferred to chilled tubes. Total protein concentration was quantified by bicinchoninic acid (BCA) assay. Lysates were adjusted to a final concentration of 5 mM DTT to prevent air oxidation. Equal amounts of total cell lysate for each sample (∼2 mg) were combined with 50 µl (100 µl of 50 % slurry) of washed FLAG-M2 magnetic beads (Sigma Aldrich) in a total volume of 1 ml made up in lysis buffer and incubated for 30 mins at 4 °C with rotation. Beads were then collected on a magnetic stand and supernatants removed. Beads were then resuspended and washed once with 1 ml lysis buffer and then four times with lysis buffer containing 500 mM NaCl. For pNPP assays, IPs were then washed in 1 ml assay buffer (50 mM Tris-HCl pH 7.4, 150 mM NaCl, 5% [v/v] glycerol, 5 mM DTT] and then resuspended in 500 µl assay buffer. Samples were then subject to pNPP assays as described above. After the assay beads were collected by magnet, washed in 1 ml TBS and resuspended in 1X SDS PAGE sample buffer and incubated at 95°C for 5 mins. Beads were collected by magnet and supernatants used for SDS-PAGE and immunoblot analysis.

### HyPer7 imaging and analysis

U2OS cells expressing PM-HyPer7 were seeded on 35 mm glass bottom IBIDI dishes (81158) prior to imaging and maintained at 37°C and 5% CO_2_. Fluorescence excitation was measured at 488 nm and 405 nm, and λem = 525 nm as described previously (*30*). Treatments were prepared at 3x and added gently to the side of the culture dish after 1 min imaging. Images were acquired using Total internal reflection fluorescence (TIRF) microscopy (Nikon Eclipse STORM-TIRF microscope), using a 40x oil immersion objective. All image analysis was conducted using ImageJ/FIJI. HyPer7 ratiometric images (λex=488/ λex=405) were false coloured using mpl-Magma look-up table (purple to yellow). GraphPad Prism (Ver. 10) was used for statistical analysis.

### HyPer7 and pervanadate reaction *in vitro*

Purified recombinant HyPer7 was buffer-exchanged into Ar-flushed 100 mM NaPOi, pH 7.4, 150 mM NaCl, 0.1 mM diethylene triamine penta-acetic acid (DTPA) using a 5 ml HiTrap desalting column (GE Healthcare). The protein concentration was determined with the Bradford reagent (Bio-Rad), using bovine serum albumin as a standard. The protein was diluted to 2.8 μM and incubated with 0.1 mM DTT for 30 min at RT. Pervanadate, H_2_O_2_ and catalase controls (subsequently referred to as “reactants”) were prepared at a 10x concentration. HyPer7 (final concentration 2.5 μM) and the reactants were mixed in a quartz cuvette, incubated for 1 min at RT and the fluorescence excitation spectrum was recorded from λex = 350 nm to 505 nm with λem = 512 nm using a Cary Eclipse Fluorescence Spectrophotometer (Agilent). The ratio of the two excitation maxima at 400 nm and 498.5 nm was the final readout. GraphPad Prism (Ver. 10) was used for statistical analysis.

### Protein kinase assays

Kinase assays were performed using nonradioactive real-time mobility shift-based microfluidic assays, as described previously (*75*), in the presence of 2 mM of the appropriate fluorescent-tagged peptide substrate (5-FAM-EGIYGVLFKKK-CONH_2_) and 1 mM ATP (unless specified otherwise). Pressure and voltage settings were adjusted manually to afford optimal separation of phosphorylated and non-phosphorylated peptides. All assays were performed in 50 mM HEPES (pH 7.4), 0.015% (v/v) Brij-35, and 5 mM MgCl_2_, using 3 ng of SRC, and real-time or end point peptide phosphorylation was determined using the ratio of the phosphopeptide:peptide. Kinase activity in the presence of different redox reagents or inhibitor compounds was quantified by monitoring the generation of phosphopeptide during the assay, relative to controls, with phosphate incorporation into the peptide generally limited to <20% to prevent ATP depletion and maintain assay linearity. The effects of variable exposure times on kinase activity were monitored by preincubating SRC and redox reagent for 10, 60 or 120 mins at room temperature (RT), and then initiating substrate phosphorylation with the addition of ATP and peptide substrate. Relative changes of SRC activity in the presence of increasing concentrations of pervanadate were monitored in real time over 4 h using 20 µM peptide substrate. The rate of substrate phosphorylation at each time point was calculated as pmol phosphate incorporation per min and plotted as a function of total assay time. Counter screen assays using 50 ng Aurora A kinase were performed as for SRC.

### Differential alkylation in cell lysates

Differential alkylation methodology was performed as previously described (*76*). HEK293T cells were plated at 2 x10^6^ per 10 cm^2^ dish until full confluency followed by PV treatments. Cells were washed twice in ice-cold PBS followed by harvesting into pH 7.0 lysis buffer [30 mM Tris-HCL (pH 7), 120 mM NaCl, 2 mM EDTA, 2 mM KCl, 10 % (v/v) glycerol, 1 % (v/v) triton, 1 % SDS (v/v), 5 mM NaF, 1X EDTA-free protease inhibitor tablet cocktail (Roche; 1836170001) and PhosSTOP tablets (Roche; 4906845001)] containing NEM (Sigma; E3876) 5 mM for alkylation. Lysates were then sonicated and incubated on ice for 20 min in the dark to enable NEM alkylation, followed by centrifugation at 13000 x g to pre-clear the lysate. During this incubation, Zeba™ spin desalting columns (Thermo Fisher Scientific; 89882) were equilibrated with lysis buffer pH 8.0 following the manufacturer’s instructions. 0.5 mg protein per sample was buffer exchanged by filtering through three Zeba™ spin columns per sample to remove excess NEM. 50 mM TCEP (Merck; 646547) was added to each sample followed by incubation at room temp for 30 mins to enable sample reduction. Samples were then filtered through three pH 8.0 lysis buffer equilibrated Zeba™ columns per sample to remove excess TCEP. Finally, samples were incubated with 10 mM EZ-Link™ Iodoacetyl-PEG2-Biotin (Thermo Fisher Scientific; 21334) for 90 mins at RT in the dark, followed by buffer exchange through three equilibrated Zeba™ spin columns to remove excess alkylating agent. Lysates were then subjected to streptavidin enrichment using the following method. Protein concentration was measured again for each sample using BCA assay and 400 µg each sample was subject to pull-down. 10 % lysate was removed and added to sample buffer for input sample. Remaining lysate was incubated with 50 µl neutravidin beads (Pierce; 29202) rotating for 2 h at 4°C. Equivalent of 10 % starting lysate concentration was removed for supernatant sample and added to sample buffer. The remaining beads were washed 4x with RIPA buffer [50 mM Tris-HCl, pH 7.4, 1% (v/v) Triton, 1% Na deoxycholate, 0.1% SDS, 150 mM NaCl, 1 mM EDTA] followed by elution in 5x sample buffer with 25 mM biotin (Merck; B4639) with incubation at 95°C for 10 mins. Input, supernatant and pull-down samples then underwent analysis by immunoblotting.

### SRC affinity precipitation from HEK293T cells

Affinity precipitation experiments used HEK293T cells transfected using a 3:1 polyethyleneimine [average Mw (weight-average molecular weight), ∼25,000 Da; Sigma-Aldrich] to DNA ratio (30:10 mg, for a single 10 cm^2^ culture dish). Proteins were harvested 48 h post transfection using a lysis buffer containing 50 mM Tris-HCl (pH 7.4), 150 mM NaCl, 0.1% (v/v) Triton X-100, and 5% (v/v) glycerol and supplemented with a protease inhibitor cocktail tablet and a phosphatase inhibitor tablet (Roche). Lysates were briefly sonicated on ice and clarified by centrifuged at 20,817g for 20 min at 4°C, and the resulting supernatants were incubated with Strep-TactinXT 4Flow affinity resin (IBA Lifesciences) for 1 h with gentle end over end mixing at 4°C. Beads containing bound protein were collected and washed three times in 50 mM tris-HCl (pH 7.4) and 500 mM NaCl and then equilibrated in storage buffer [50 mM Tris-HCl (pH 7.4), 100 mM NaCl, and 5% (v/v) glycerol]. The purified kinases were then proteolytically eluted from the beads over a 1 h period using 3C protease (0.5 mg) at 4°C with gentle agitation and then assayed using real-time microfluidic peptide assay as described above. For MS-dependent detection of cysteine modifications, HEK293T cells were treated with pervanadate or H_2_O_2_ for 30 mins, and protein harvested in 50 mM Tris-HCl (pH 6.5), 150 mM NaCl, 0.1% (v/v) Triton X-100, and 5% (v/v) glycerol supplemented with a protease and phosphatase inhibitor tablets (Roche), and 100 mM iodoacetamide to alkylate thiols. After brief sonication, lysates were incubated in the dark for 1 h at 4°C, clarified by centrifugation and affinity purified as above.

### Identification of affinity enriched SRC cysteine oxidation state using mass spectrometry

Affinity purified SRC preparations were diluted to 180 μl in 100 mM ammonium bicarbonate pH 8.0 and reduced and alkylated with dithiothreitol and iodoacetamide, as previously described by (*77*). Samples were subject to SP3-bead digestion as described by (*78*) with the adaptions of: using 100 mM ammonium bicarbonate pH 8.0, 0.5 μg Trypsin gold (Promega) and post-digest bead washing in 1% (w/v) Rapigest (Waters). Once supernatants were combined, samples were acidified by addition of a final concentration of 0.5% TFA and incubated for 30 min each at 37 °C with 600 rpm shaking and on ice. Samples were centrifuged at 13000 x g for 10 min at 4 °C and cleared supernatant collected into fresh tubes prior to vacuum centrifugation. Dried peptides were solubilized in 20 μl of 3% (v/v) acetonitrile and 0.1% (v/v) TFA in water, sonicated for 10 minutes, and centrifuged at 13000 x g for 15 min at 4 °C prior to reversed-phase HPLC separation using an Ultimate3000 nano system (Dionex) over a 60 min gradient, as described by (*79*). All data acquisition was performed using a Thermo QExactive HF mass spectrometer (Thermo Fisher Scientific), with higher-energy C-trap dissociation (HCD) fragmentation set at 29% normalized collision energy for 2+ to 5+ charge states. MS1 spectra were acquired in the Orbitrap (60K resolution at 200 m/z) over a m/z range of 350 to 2000, AGC target = 3e6, maximum injection time = 100 ms, with an intensity threshold for fragmentation of 1e3. MS2 spectra were acquired in the Orbitrap (30K resolution at 200 m/z), maximum injection time = 45 ms, AGC target = 1e5 with a 20 s dynamic exclusion window applied with a 10-ppm tolerance. Data was analyzed using Proteome Discoverer 2.4 (Thermo Fisher Scientific) in conjunction with MASCOT (*80*); searching the UniProt Human Reviewed database (updated weekly, accessed March 2022) with constant modifications = carbamidomethyl (C), variable modifications = oxidation (M), phosphorylation (ST), trioxidation (C), instrument type = electrospray ionization–Fourier-transform ion cyclotron resonance (ESI-FTICR), MS1 mass tolerance = 10 ppm, MS2 mass tolerance = 0.01 Da with the ptmRS node active. Label free quantification was performed using the Minora feature detector node, calculating the area under the curve for m/z values, total protein abundance was determined using the HI3 method (*81*). For cysteine trioxidation-containing peptides identified in at least 3 replicates, peptide abundance was normalized against the calculated total protein abundance to account for potential protein load variability during analysis prior to normalization against untreated control. For the best MASCOT scoring PSM of a C188-trioxidation containing peptide, the spectrum was extracted in *.mgf format from the raw file and imported into a custom R script for re-drawing and manual annotation.

### Differential scanning fluorimetry

Thermal shift assays were performed using a StepOnePlus real-time polymerase chain reaction (PCR) machine (Life Technologies) using SYPRO Orange dye (Invitrogen) and thermal ramping (0.3°C in step intervals between 25°C and 94°C). Recombinant SRC proteins were assayed at a final concentration of 2.5 mM in 50 mM tris-HCl (pH 7.4) and 100 mM NaCl in the presence or absence of the indicated concentrations of ATP, H_2_O_2_, DTT, orthovanadate, pervanadate or Dasatinib [final dimethyl sulfoxide (DMSO) concentration no higher than 4% (v/v)] as described previously (*75*). Normalized data were processed using the Boltzmann equation to generate sigmoidal denaturation curves, and average Tm/ΔTm values were calculated as previously described (*82*) using GraphPad Prism software.

### Hydrogen-Deuterium exchange mass spectrometry

A preincubation was performed by adding to a concentrated SRC sample (57 μM SRC in 50 mM Tris pH 7.5, 150 mM NaCl) a 4.5 mM PV solution to yield a final solution of 5 μM SRC (20 mM Hepes pH 7, 150 mM NaCl) in presence of 0.5 mM or 2.5 mM of PV. Samples were incubated for 15 minutes at room temperature and then exposed for different deuteration times (3 seconds on ice, 3, 30, 300 and 3000 seconds at room temperature) to a deuterated buffer: 50 mM Tris pH 7.5, 150 mM NaCl, with D_2_O at a final concentration of 90% (Thermo Fisher Scientific, 166300100) to yield a final solution that was 0.5 μM of SRC in absence/presence of 0.05 mM or 0.25 mM of PV. Triplicate samples were aliquoted for each deuteration timepoint. Exchange reactions were quenched in 2 M Guanidine-Hydrochloride, 100 mM Glycine, formic acid (Fisher Chemical, 10596814), LC-MS grade water (Romil, H949), pH 2.3 (final pH was 2.5). Samples were then flash-frozen in liquid nitrogen immediately and stored at -80 °C until analysis. Prior to LC-MS analysis, samples were quickly thawed and injected on a HDX sample manager coupled to an Acquity ultra performance liquid chromatography (UPLC) M-Class system (Waters) set at 0.1 °C. Samples were then digested at 15 °C using a 20 mm x 2.0 mm home-made pepsin column (Pepsin from Thermo Scientific, 20343 and hardware from Upchurch Scientific, C130-B) and loaded on a UPLC pre-column (Acquity UPLC BEH C18 VanGuard pre-column, 2.1 mm I.D. × 5 mm, 1.7 μm particle diameter, Waters, 186003975) for 2 min at 100 μL/min. Digested peptides were then eluted from the pre-column onto an Acquity UPLC BEH C18 column (1.0 mm I.D. × 100 mm, 1.7 μm particle diameter, Waters, 186002346) using a 5-43% gradient of Acetonitrile (Romil, H050), 0.1% formic acid over 12 min. MS data acquisition was done on a Synapt G2Si-HDMS (Waters) with positive electrospray ionization, using HDMS^E^ acquisition mode over a *m*/*z* range of 50–2000 and a 100 fmol.μL^-1^ Glu-FibrinoPeptide solution was used for lock-mass correction and calibration. The following parameters were used during the acquisition: capillary voltage, 3 kV; sampling cone voltage, 40 V; source temperature, 90°C; desolvation gas, 150°C and 650 L.h^−1^; scan time, 0.3 s; trap collision energy ramp, 20–45 eV. Peptide identification was performed using ProteinLynx Global Server 3.0.1 (Waters) with a home-made protein sequence library containing SRC and pepsin sequences, with peptide and fragment tolerances set automatically by PLGS. Deuterium uptakes for all identified peptides were then filtered and validated manually using DynamX 3.0 (Waters) as follows: only peptides identified with a minimum fragment per amino acid of 0.2, a minimum intensity of 10^3^, a length between 5 and 25 residues, a minimum PLGS score of 6.62 and a MH^+^ tolerance of 5 ppm were kept. An initial automated spectral processing step was conducted by DynamX followed by a manual inspection of individual peptides for sufficient quality where only one charge state per peptide was kept. Deuterium uptake is not corrected for back-exchange and is reported as relative values. HDX-MS results were statistically validated using an in-house program (Archaeopteryx), where the t-statistic threshold was set to 0.05. Only statistically significant peptides showing a difference greater than 0.25 Da and 2% for at least two timepoints were kept. HDX-MS results were illustrated on SRC open and closed conformations (PDB: 2SRC and 1Y57 respectively) using PyMOL 2.5.4 (www.pymol.org).

### AlphaFold3 modeling

SRC SH2 and C-terminal pTyr complexes were modelled from sequence using AlphaFold3 through its public web server (https://www.alphafoldserver.com). Standard AlphaFold procedures were followed, and the outputs were manually inspected to confirm the correct orientation of the SH2 domain bound to peptide. Post-translational modifications were added using the PTMs function on the selected residues. For each complex, five models were generated, and their confidence metrics were evaluated using the AlphaFold3 ranking scores. The highest-ranking model, determined by its pLDDT score, was selected for further analysis. Structural representations were prepared using UCSF Chimera, and alignment was performed using MatchMaker in Chimera X.

### Isothermal Titration Calorimetry

ITC experiment was measured by MicroCal PEAQ-ITC Automated at 25℃. After treatment, SRC SH2 domains and peptides were buffer exchanged into fresh experimental buffer using ultrafiltration prior to titrations. ITC experiments were performed by stepwise titration of the peptides (400 μM) into an adiabatic cell containing SRC SH2 (40 μM), and the heat energy change accompanying the reaction was detected upon each injection by comparison with a reference cell. The stoichiometry of binding (n) was between 0.84 and 1.1 for all experiments. Protein solution was placed in the 380 μl calorimeter cell and stirred at 750 rpm to ensure rapid mixing, and 2 μl aliquots of the peptides were injected with a 120 s interval between each injection until saturation. The heat changes were integrated after subtracting values obtained when peptides were titrated into buffer, data were analyzed with the MicroCal PEAQ-ITC software.

### Cell counting and Incucyte® imaging

MEFs were plated on fibronectin (Roche #10838039001) 5 µg/cm^2^ coated 6-well plates at 0.2 x10^6^ and grown over 6 days with a media change on day 3. On daily intervals, one well was trypsinized, resuspended in normal growth medium and counted with trypan blue using the countess cell counter (Invitrogen) for the manual growth curves. For live imaging, MEFs were plated on fibronectin (Roche; 10838039001) 5 µg/cm^2^ coated CytoOne 96-well plates at 2.5 x10^5^ cells per well. The following day, the plate was transferred to an Incucyte^®^ S3 live cell imaging and analysis system (Sartorious; IC70682) and 4 regions were imaged per well every 2 h for 6 days. Media was refreshed on day 3.

### Immunofluorescence microscopy

Cells were plated on fibronectin (Roche #10838039001) 5 µg/cm^2^ coated coverslips at 0.2 x10^6^ and let to grow for 6 days with a media change on day 3. Then fixed using 4% paraformaldehyde in PBS for 10 min. The cells were then permeabilized in 0.5% Triton X-100 and 3% BSA in PBS for 2 min and blocked in 0.2% Triton X-100 and 3% BSA in PBS (blocking buffer) for 1 h. They were stained using Alexa Fluor™ Plus 647 Phalloidin (1:200) and Hoechst (1:2000) for 1 h then washed 3 times in blocking buffer for 1 h. Images were taken using and Leica Stellaris 8 with a 40X objective and 1.5 zoom. The nucleus of 6 different fields per condition were counted.

## Ethical Statement

All animal experiments at The Babraham Institute were reviewed and approved by The Animal Welfare and Ethics Review Body and performed under Home Office Project license PP4624082.

## Supplementary materials

Figs. S1 to S6

Movies S1-S6

Table S1.

## Acknowledgements

We thank the Babraham Institute imaging facility, J. Clark in the Babraham Institute biological chemistry facility, C. Switzer for discussions about redox chemistry, T. Chessa for advice on retroviral transduction, K. Wojdyla for advice on differential alkylation, and E. Veal, P. Hawkins, A. Tuncay, R. Williams and members of the Sharpe lab for critical feedback on the manuscript.

## Author contributions

Conceptualization: KEM, DPB, HJS. Methodology: KEM, MB, NC, OR, DE, LAD, SAC, DPB. Investigation: KEM, NC, OR, DE, DPB. Formal analysis: KEM, MB, NC, OR, DE, LAD, SAC, DPB. Visualization: KEM, MB, NC, OR, DE, TL, SAC, DPB, HJS. Validation: KEM, DPB. Data curation: HEJ, HJS. Funding acquisition: PC, CEE, JM, PAE, HJS. Project Administration: HJS. Supervision: PC, CEE, JM, PAE, HJS. Resources: PC, CEE, JM, PAE, HJS. Writing – original draft: KEM, DPB, HJS. Writing – review and editing: KEM, MB, NC, CEE, JM, PAE, DPB, HJS.

## Data and materials availability

Mass spectrometry data have been deposited to the ProteomeXchange Consortium through the PRIDE partner repository with the dataset identifiers PXD059503 (CPT enrichment), PXD059641(Cysteine S-sulfonate modifications) and PXD059840 (HDX-MS). All other materials are available upon request to the corresponding authors.

