## Supplementary Material for "Pervanadate-induced oxidation relieves autoinhibition of SRC protein tyrosine kinase"

**Supplementary Figures and legends**

Figure S1

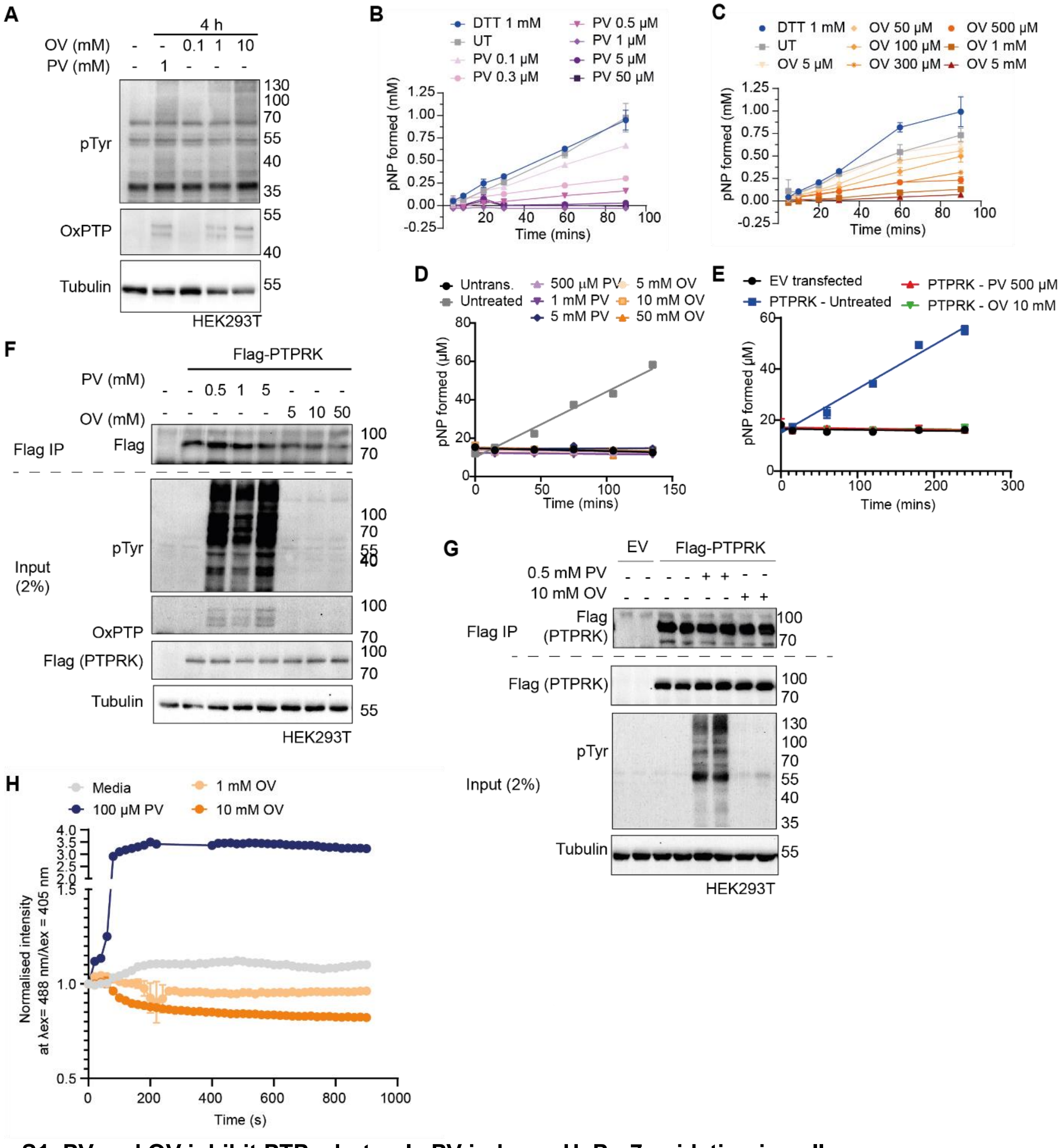


**Fig. S1. PV and OV inhibit PTPs, but only PV induces HyPer7 oxidation in cells.**
(**A**) HEK293T cells were treated as shown, lyzed and subjected to immunoblot analysis with the indicated antibodies.
(**B-C**) pNPP phosphatase assays with 0.5 μM recombinant PTPRK intracellular domains with the indicated concentrations of PV, OV or DTT. UT = untreated.
(**D**) Immunoprecipitates analyzed in F were subjected to a pNPP colorimetric phosphatase assay.
(**E**) Immunoprecipitates analyzed in G were subjected to a pNPP colorimetric phosphatase assay. EV= empty vector.

(**F**) HEK293T cells were transiently transfected with empty vector or a plasmid for expression of flag-tagged PTPRK intracellular domain and treated as shown. Cells were lysed and PTPRK was flag immunoprecipitated. Inputs and immunoprecipitates were subject to immunoblot analysis using the indicated antibodies. Image representative of N=2 experiments.
(**G**) As for F.
(**H**) Quantification of the ratio of the two excitation maxima (F488/405), normalized against initial value, of PM-HyPer7 expressing U2OS cells over a time course, imaged every 20 s. Treatments were added at 60 seconds (arrow). The bars represent mean of n=3 ± SD. ** $p \leq 0.001$ Unpaired t-test at t= 400 s.

### Figure S2

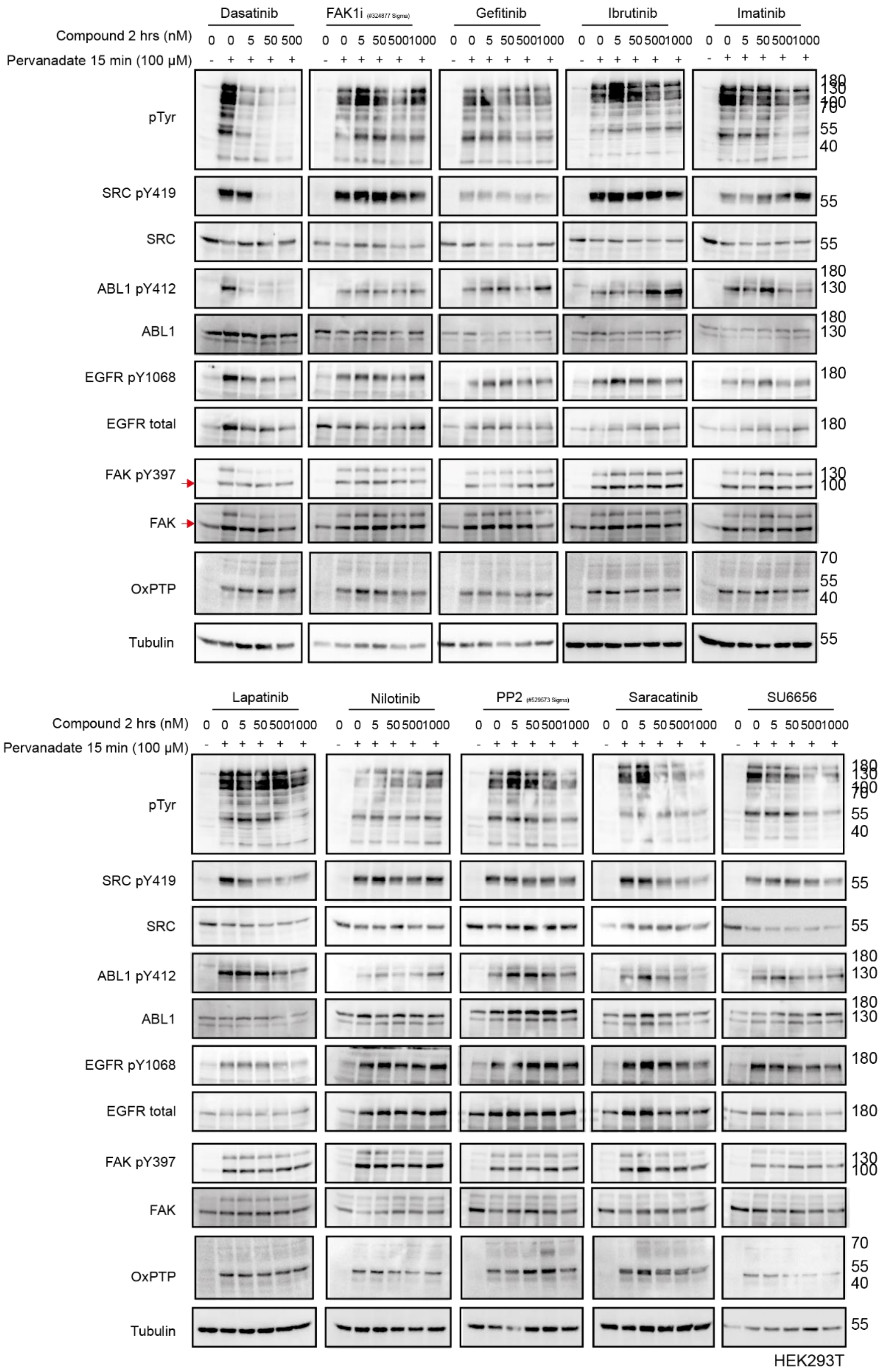
Dasatinib
FAK1i
Gefitinib
Ibrutinib
Imatinib
Compound 2 hrs (nM)
Pervanadate 15 min (100 µM)
pTyr
SRC pY419
SRC
ABL1 pY412
ABL1
EGFR pY1068
EGFR total
FAK pY397
FAK
OxPTP
Tubulin
Lapatinib
Nilotinib
PP2
Saracatinib
SU6656
HEK293T

**Fig. S2. SFKs are important for PV-driven pTyr accumulation.** HEK293T cells were pretreated with the indicated tyrosine kinase inhibitors and concentrations for 2 hours with or without PV (15 minutes) before lysis and immunoblotting with antibodies shown.

Figure S3

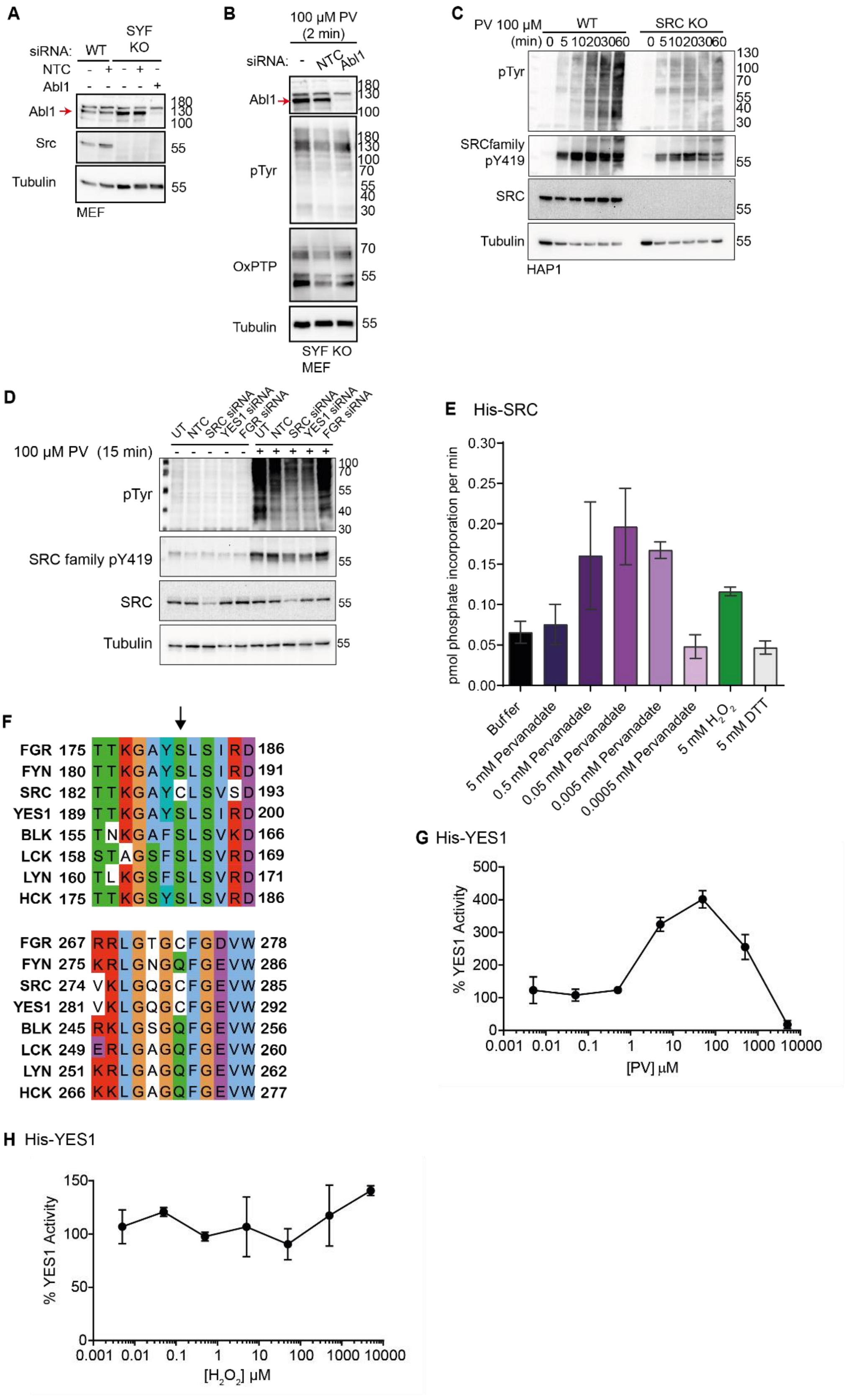

A
siRNA: WT SYF KO
NTC
Abl1
Abl1
Src
Tubulin
MEF
B
100 µM PV (2 min)
siRNA: - NTC Abl1
Abl1
pTyr
OxPTP
Tubulin
SYF KO MEF
C
PV 100 µM WT SRC KO
(min) 0 5 10 20 30 60 0 5 10 20 30 60
pTyr
SRCfamily pY419
SRC
Tubulin
HAP1
D
UT NTC SRC siRNA YES1 siRNA FGR siRNA UT NTC SRC siRNA YES1 siRNA FGR siRNA
100 µM PV (15 min)
pTyr
SRC family pY419
SRC
Tubulin
E His-SRC
pmol phosphate incorporation per min
Buffer
5 mM Pervanadate
0.5 mM Pervanadate
0.05 mM Pervanadate
0.005 mM Pervanadate
0.0005 mM Pervanadate
5 mM H2O2
5 mM DTT
F
FGR 175 TTKGAYSLSIRD 186
FYN 180 TTKGAYSLSIRD 191
SRC 182 TTKGAYCLSVSD 193
YES1 189 TTKGAYSLSIRD 200
BLK 155 TNKGAFSLSVKD 166
LCK 158 STAGSFSLSVRD 169
LYN 160 TLKGSFSLSVRD 171
HCK 175 TTKGSYSLSVRD 186
FGR 267 RRLGTGCFGDVW 278
FYN 275 KRLGNGQFGEVW 286
SRC 274 VKLGQGCFGEVW 285
YES1 281 VKLGQGCFGEVW 292
BLK 245 RKLGSGQFGEVW 256
LCK 249 ERLGAGQFGEVW 260
LYN 251 KRLGAGQFGEVW 262
HCK 266 KKLGAGQFGEVW 277
G His-YES1
% YES1 Activity
[PV] µM
H His-YES1
% YES1 Activity
[H2O2] µM

**Fig. S3. PV and $H_2O_2$ dose-dependently activate SRC.**

(**A**) WT and Src/Yes/Fyn (SYF) KO cells were transfected with or without NTC or ABL1 siRNAs, prior to lysis and immunoblotting with indicated antibodies. Red arrow indicates the band for ABL1.

(**B**) Src/Yes/Fyn (SYF) KO cells were transfected with or without NTC or ABL1 siRNAs andtreated with 100 µM PV for 2 min, prior to lysis and immunoblotting with indicated antibodies. Red arrow indicates the band for ABL1.

(**C**) Hap1 WT and SRC KO cells were treated with 100 µM PV for indicated times prior to lysis and immunoblotting with antibodies shown.

(**D**) HEK293T cells were transfected with or without NTC or SFK siRNAs and treated with or without 100 µM PV for 15 min, followed by a washout and incubation with indicated concentrations of GSH, with or without pre-incubation.

(**E**) Change in the rate of SRC dependent substrate activity over time (adapted from Fig. S4A). The relative rate of phosphate incorporation (pmol per min) was calculated at each time point and is plotted as a function of time. Means ± SD of three experimental repeats are shown.

(**F**) Alignment of SFKs highlighting the amino acids surrounding the equivalent of SRC Cys 188 (upper) and Cys 280 (lower). Alignment generated using ClustalOmega (*83*) and edited in Jalview (84) with Clustal color scheme.

(**G**) YES1 (40 ng) dependent phosphorylation of a specific fluorescent substrate (2 µM) measured in real time in the presence of the indicated concentrations of pervanadate following pre-incubation for 10 min. Reactions were initiated with the addition of peptide substrate and ATP, and the rate of peptide phosphorylation was measured after 15- and 25-min assay time. Data is % YES1 activity relative to a buffer only control, and means ± SD of three experimental repeats are shown.

(**H**) Dose response curve for $H_2O_2$ pre-incubated with recombinant 6xHis-YES1 (40 ng) for the indicated time periods. Reactions were initiated with the addition of peptide substrate and ATP, and phosphorylation was measured after 15- and 25-min assay time. Means ± SD of three experimental repeats are shown.

Figure S4

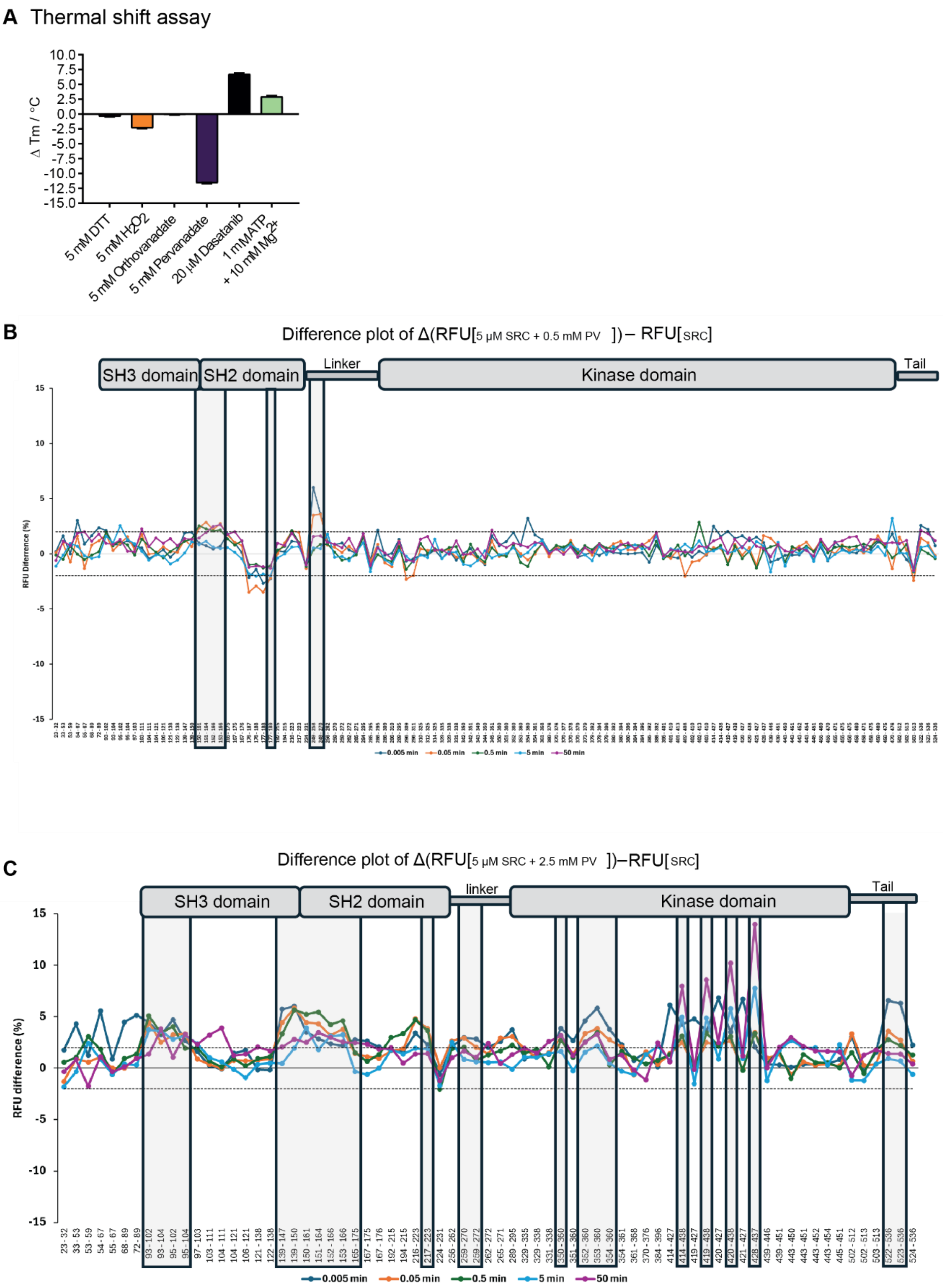
A Thermal shift assay
Δ Tm / °C
5 mM DTT
5 mM H2O2
5 mM Orthovanadate
5 mM Pervanadate
20 μM Dasatanib
1 mM ATP + 10 mM Mg2+
B
Difference plot of Δ(RFU[5 μM SRC + 0.5 mM PV ])– RFU[SRC]
SH3 domain
SH2 domain
Linker
Kinase domain
Tail
RFU Diferrence (%)
0.005 min
0.05 min
0.5 min
5 min
50 min
C
Difference plot of Δ(RFU[5 μM SRC + 2.5 mM PV ])–RFU[SRC]
SH3 domain
SH2 domain
linker
Kinase domain
Tail
RFU difference (%)
0.005 min
0.05 min
0.5 min
5 min
50 min

**Fig. S4. PV induces differential relative fractional uptake across SRC peptides.**
(**A**) Thermal shift analysis of 2.5 μM purified full-length SRC in the presence of 5 mM PV, OV, DTT, and $H_2O_2$, 1 mM ATP + 10 mM $MgCl_2$, and 20 μM Dasatinib. Means of n=2 ± SD are shown.
(**B**) Relative fractional uptake (RFU) difference plots are represented for HDX-MS analysis of SRC for 0.5 mM PV versus untreated control. RFU differences are depicted for 0.005, 0.05, 0.5, 5 and 50 minutes of deuteration. Framed peptides represent the identified peptides for which the difference of deuterium incorporation has been statistically validated for at least two timepoints.
(**C**) Relative fractional uptake difference plots are represented for HDX-MS analysis of SRC for 0.5 mM PV versus untreated control. RFU differences are depicted for 0.005, 0.05, 0.5, 5 and 50 minutes of deuteration. Framed peptides represent the identified peptides for which the difference of deuterium incorporation has been statistically validated for at least two timepoints.

Figure S5

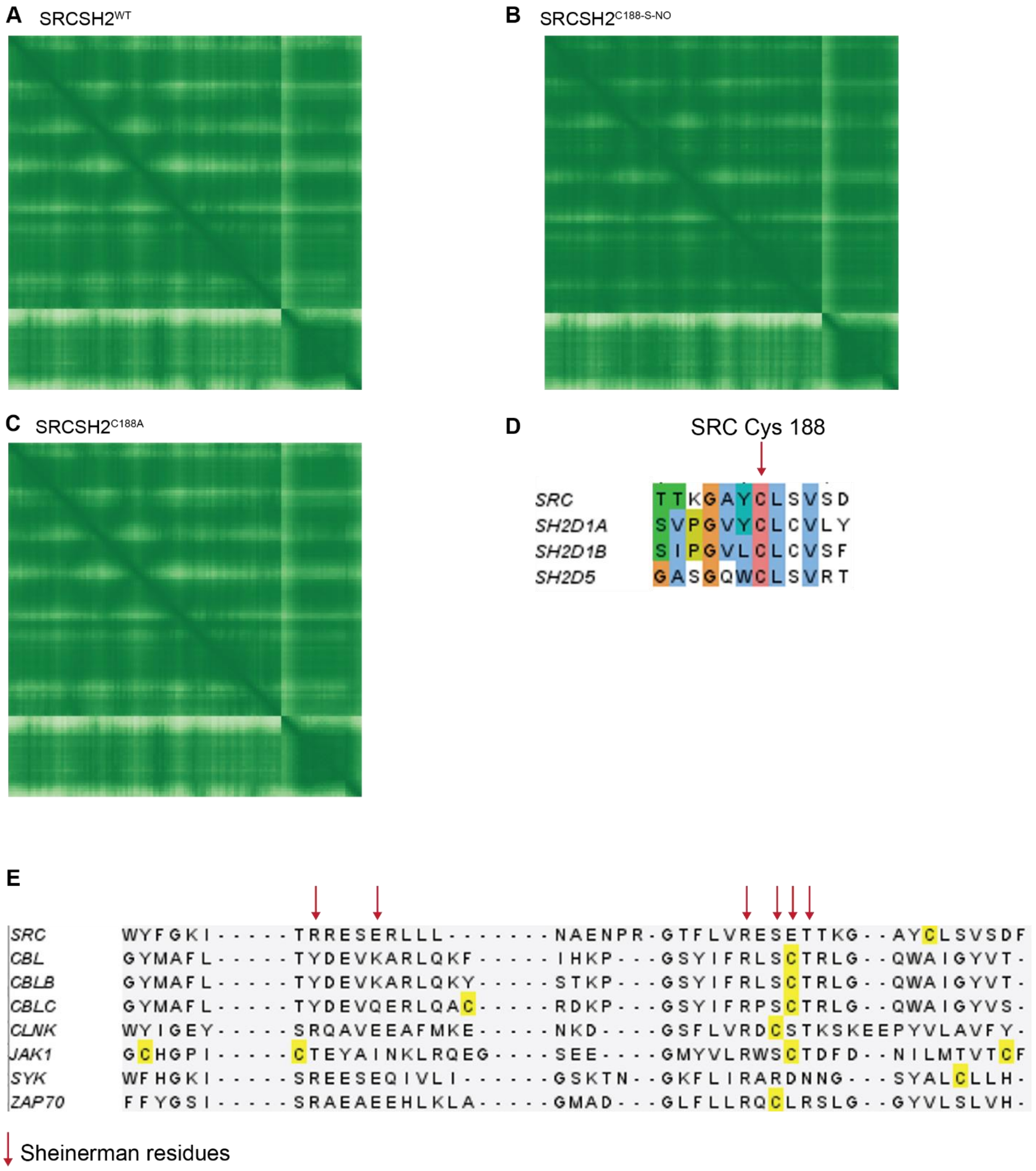


**Fig. S5. AlphaFold3 (AF3) error plots corresponding to models shown in Fig. 4C.**
(**A**) AF3 error plot for SRC-SH2 interaction with SRC C terminal phosphopeptide.
(**B**) As for A but for SRC-SH2$^{C188\text{-}S\text{-}NO}$.
(**C**) As for A but for SRC-SH2$^{C188A}$.
(**D**) Clustal alignment of region spanning SRC Cys 188 with equivalent regions in the SH2 domains of SH2D1A, SH2D1B, SH2D5. Edited in Jalview (*2*).
(**E**) Alignment highlighting cysteine residues within the pTyr binding pocket of SH2 domains from indicated proteins. Alignments derived from SH2DB (85). Edited in Jalview. Sheinerman residues highlighted by red arrow.

Figure S6

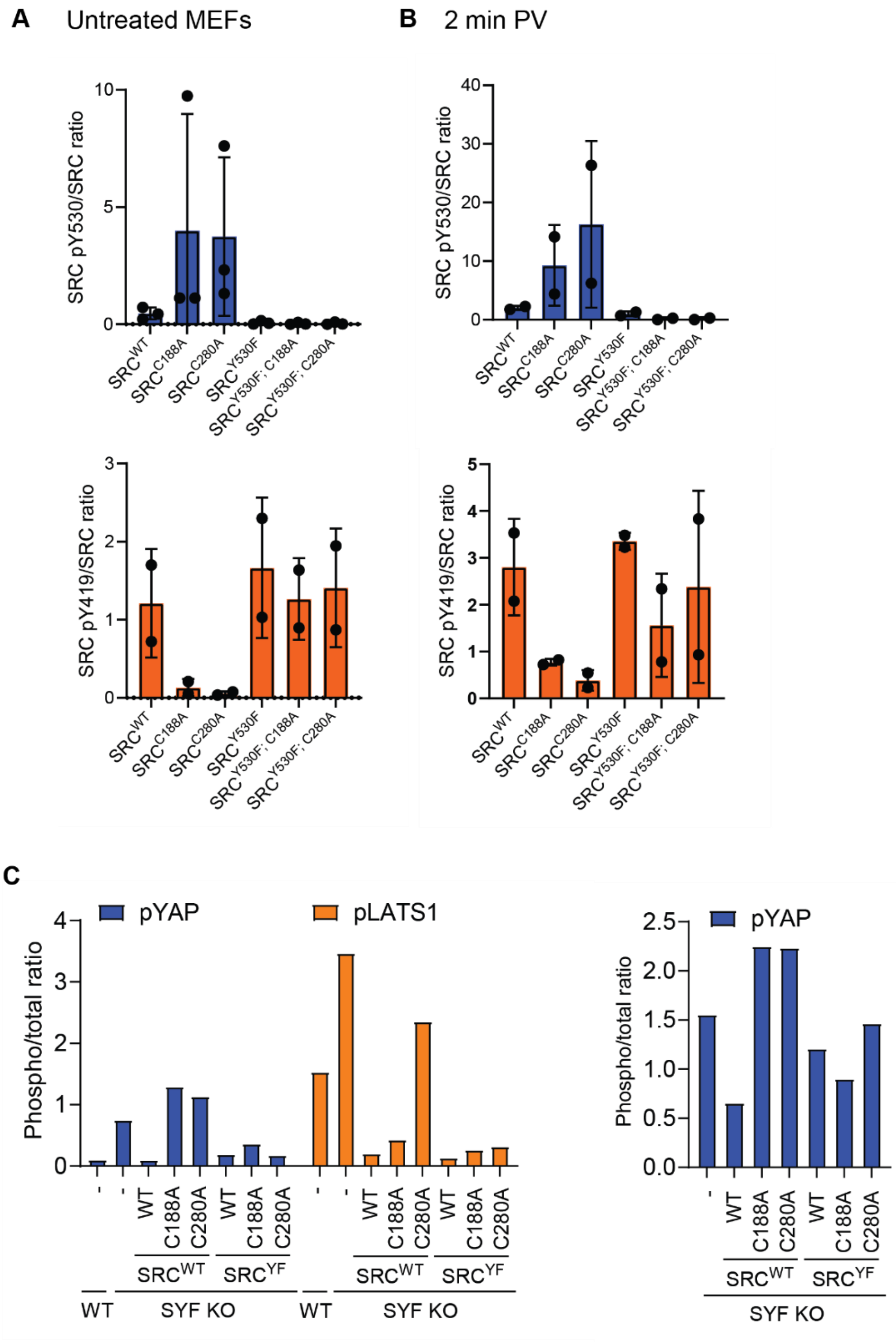


**Fig. S6. SRC cysteines are required to overcome contact inhibition of proliferation**

(**A**) Densitometric quantification of Western blot in Fig. 5A and replicate experiments. The ratio of phosphorylated/total levels are shown for n=3 for pY530 and n=2 for pY419. Means ± SD are shown.

(**B**) Densitometric quantification of 2 min PV treated samples from Western blot in fig. 5A and replicate experiment. The ratio of phosphorylated/total levels are shown for n=2 for pY530 and n=2 for pY419. Means ± SD are shown.

(**C**) Densitometric quantification of Western blot shown in Fig. 5E (left), and a replicate blot (right).

**Other supplementary files:**

Movie S1. TIRF microscopy of U2OS cells expressing PM-HyPer7 treated with media. Related to Fig. 1.

Movie S2. TIRF microscopy of U2OS cells expressing PM-HyPer7 treated with 100 μM PV. Related to Fig. 1.

Movie S3. Brightfield timelapse of SYF KO MEF cells in culture. Related to Fig. 5.

Movie S4. Brightfield timelapse of SYF KO + SRC MEF cells in culture. Related to Fig. 5.

Movie S5. Brightfield timelapse of SYF KO + $SRC^{C188A}$ MEF cells in culture. Related to Fig. 5.

Movie S6. Brightfield timelapse of SYF KO + $SRC^{C280A}$ MEF cells in culture. Related to Fig. 5.

Table S1. Mass spectrometry data for cysteine oxidation of affinity purified SRC.
